# Strategic control of location and ordinal context in visual working memory

**DOI:** 10.1101/2022.12.10.519889

**Authors:** Jacqueline M. Fulvio, Qing Yu, Bradley R. Postle

## Abstract

Working memory (WM) requires encoding stimulus identity and context (e.g., where or when stimuli were encountered). To explore the neural bases of the strategic control of context binding in WM, we acquired fMRI while subjects performed delayed recognition of three orientation patches presented serially and at different locations. The recognition probe was an orientation patch with a superimposed digit, and pretrial instructions directed subjects to respond according to its location (*location-relevant*), to the ordinal position corresponding to its digit (*order-relevant*), or to just its orientation (relative to all three samples; *context-irrelevant*). Delay-period signal in PPC was greater for context-relevant than for *context-irrelevant* trials, and multivariate decoding revealed strong sensitivity to context binding requirements (relevant vs. *irrelevant*) and to context domain (*location-* vs. *order-relevant*) in both occipital cortex and PPC. At recognition, multivariate inverted encoding modeling revealed markedly different patterns in these two regions, suggesting different context-processing functions. In occipital cortex, an active representation of the location of each of the three samples was reinstated, regardless of trial type. The pattern in PPC, in contrast, suggested a trial type-dependent filtering of sample information. These results indicate that PPC exerts strategic control over the representation of stimulus context in visual WM.

## Introduction

Several recent studies have provided evidence that delay-period activity in the intraparietal sulcus (IPS) reflects, at least to some extent, context-binding operations (Cai et al. 2019; Cai et al. 2020; Gosseries, Yu, et al. 2018), thus offering a complement to the idea that this activity reflects stimulus representation in visual working memory (VWM; Bettencourt & Xu 2016; Todd & Marois 2004; Xu 2017; Xu & Chun 2006). Gosseries, Yu, et al. (2018) sought to dissociate activity related to context-binding demands from that related to memory load, per se, by varying stimulus category homogeneity within the memory set. In addition to trials requiring delayed recall of the direction of motion of one dot-motion patch (“*1M*”), subjects also performed two types of load-of-three trials: memory for three motion patches (“*3M*”); and memory for one motion patch and two color patches (“*1M2C*”). Items were presented serially, and a digit in the middle of the response dial indicated which sample (the first, second, or third) was to be recalled. Thus, remembering ordinal context was critical for *3M* trials, but much less so for *1M2C* trials, for which the ordinal position of only one of the two color patches needed to be retained. Delay-period activity in IPS was elevated for *3M* trials relative to *1M2C* and *1M* trials, which themselves did not differ.

Cai et al. (2020) used a logic similar to that of Gosseries, Yu, et al. (2018), but used location as the critical dimension of context instead of ordinal position: sample items could appear in four possible locations, and trials required delayed recall of one oriented bar (“*1O*”), of one from a set of three simultaneously presented oriented bars (“*3O*”), or of one item from a set of one orientated bar, one color patch, and one luminance patch (“*1O1C1L*”). For all trial types, the location at which the response dial appeared matched the location of the sample item to be recalled. However, because the orientation, color, and luminance response dials only afforded a response to one kind of stimulus domain, the context-binding demands on *1O1C1L* trials were negligible. As was the case with Gosseries, Yu, et al. (2018), delay-period activity in IPS was markedly higher for trials with high context-binding demand (i.e., *3O*) relative to *1O* and to *1O1C1L* trials, which did not differ (Cai et al. 2020).

The above-summarized studies suggest an alternative interpretation to the pattern of load-sensitivity that is routinely observed in IPS: Although it has traditionally been interpreted as evidence for a role for IPS as a VWM buffer (Bettencourt & Xu 2016; Todd & Marois 2004; Xu 2017; Xu & Chun 2006), it might reflect, at least in part, a role for IPS in context binding. Because Gosseries, Yu, et al. (2018) only assessed ordinal position, and (Cai et al. 2020) only spatial location, an important question to address is whether the same areas of IPS are involved in the processing of context from both of these domains. A second important question is whether context-binding in VWM can be strategically controlled according to task demands. An alternative, implied in results from a different set of analyses not reviewed here (Cai et al. 2019), raises the possibility that location context may be encoded obligatorily into WM, even when it is task irrelevant.

The current study addressed three outstanding questions about context binding in visual working memory. Empirically, because the designs of Gosseries, Yu, et al. (2018) and of Cai et al. (2020) confounded context-binding demands with stimulus type, it would seek evidence for selective sensitivity of IPS to the manipulation of context binding, above and beyond its sensitivity to load, in a task in which the stimulus content (three orientation patches) and presentation (serial presentation at three different locations) were identical across conditions, and a pre-trial instructional cue indicated whether location context, ordinal context, or neither was required to interpret the memory probe. Theoretically, there were two key questions. The first was to explore how the brain processes context differently as a function of its informational domain (here, location versus order). (I.e., although Gosseries, Yu, et al. (2018) documented IPS (and frontal) sensitivity to the manipulation of ordinal context, and Cai et al. (2020) documented IPS sensitivity to the manipulation of location context, the processing of context in each of these domains has not been compared directly.) The second theoretical question was whether the processing of stimulus context is under strategic control. (E.g., location context would vary in the same way on *location-relevant* and *order-relevant* trials, but would its processing differ as a function of its relevance for behavior?) The study design was preregistered (https://osf.io/gc9m4/?view_only=c243e13b59294a06bd299b6d06c63a1c) after functional magnetic resonance imaging (fMRI) scanning of three pilot subjects confirmed that our design was practical, and the preregistration plan was organized into three hypotheses corresponding to these three questions. For completeness and transparency, the preregistered hypotheses are presented in Table 1. For clarity of exposition, however, the Methods and Results sections are organized by question: context binding vs. load; domain specificity of context binding; and controllability of context binding.

**Table 1.** Preregistered hypotheses.

|  |  |
| --- | --- |
| *Hypothesis 1 (context binding vs. load; assessed with univariate analyses)* |  |
| *Hypothesis 1A:* | Late delay-period fMRI signal intensity (at 10-14s; TRs 6-7) in the *parietal-delay* ROI will be modulated by context-binding demands, with greater signal intensity for context-relevant trials compared *context-irrelevant* and load-of-1 trials. |
| *Hypothesis 1B:* | Late delay-period fMRI signal intensity in the *occipital-sample* ROI will not be modulated by context-binding demands, with signal returning to baseline in all four trial types. |
| *Hypothesis 1C:* | Late delay-period fMRI signal intensity will not differ between location-context and ordinal-context trials, in either the *parietal-delay* ROI or the *occipital-sample* ROI. |
| *Hypothesis 2 (domain specificity of context processing; assessed with multivariate pattern analyses)* |  |
| *Hypothesis 2A:* | Multivariate pattern analysis (MVPA) of activity in the *parietal-delay* ROI will be successful in discriminating context-relevant from *context-irrelevant* conditions (i.e., [(*location-relevant* + *order-relevant*) versus *context-irrelevant*] trials) during all three epochs of the trial. |
| *Hypothesis 2B:* | MVPA of activity in the *occipital-sample* ROI will be successful in discriminating context-relevant from *context-irrelevant* conditions (i.e., [(*location-relevant* + *order-relevant*) versus *context-irrelevant*] trials) during all three epochs of the trial. |
| *Hypothesis 2C:* | MVPA of activity in the *parietal-delay* ROI will be successful in discriminating *location-relevant* from *order-relevant* trials during all three epochs of the trial. |
| *Hypothesis* | MVPA of activity in the *occipital-sample* ROI will be successful in discriminating |
| 2D: | location-relevant from order-relevant trials during all three epochs of the trial. |
| *Hypothesis 3 (strategic control of context processing; assessed with inverted encoding modeling)* |  |
| *Hypothesis*<br>3A: | Multivariate inverted encoding modeling (IEM) of activity in the *occipital-sample* ROI will produce robust reconstructions of the probe location (TRs 9 and 10) from load-of-3 trials when trained on data corresponding to the sample location-evoked signal (TR 4) of load-of-1 trials, for all load-of-3 trial types. |
| *Hypothesis*<br>3B: | At probe (TRs 9 and 10), IEM reconstruction of the location of the sample corresponding to the digit will be significantly different from the IEM reconstruction of the location of the probe in the *occipital-sample* ROI activity. |
| *Hypothesis*<br>3C: | At probe (TRs 9 and 10), IEM reconstruction of the location of the uncued sample will be significantly different from the IEM reconstruction of the location of the probe in the *occipital-sample* ROI activity. |

## Materials and Methods

### Subjects

Estimated effect sizes for the preregistered study were based on data from previous experiments by our group (Gosseries, Yu, et al. 2018; Cai et al. 2020; Yu, Teng, & Postle 2020). Power analyses based on the results of those studies indicated that we would need data from 15 subjects to achieve 90% power to detect the effects predicted by Hypotheses 1-3. Subjects who met the following inclusion criteria were enrolled in the order in which they volunteered: being 18-35 years in age, right-handed, having normal or corrected-to-normal vision, reporting no history of neurological disease, seizures, or fainting, of a history chronic alcohol consumption or of psychotropic drugs, and having no contraindications for MRI scanning. Subjects who were able to achieve an accuracy of 83% correct or higher in each of the three context-binding conditions (see *Load-of-3 trials*, below, for details) for at least one of six 18-item training blocks administered during a behavioral training/screening session were invited to continue in the fMRI portion of the study. Subjects with fMRI datasets deemed unusable were replaced, and data collection continued until 15 usable datasets were obtained.

34 individuals completed the behavioral training/screening session with the performance of 23 of those individuals qualifying for the fMRI portion of the study. 20 total were scanned, with data from 5 subjects deemed unuseable due to excessive errors and/or missed responses (n=2), completion of only one of the two scanning sessions (n=2), or findings of clinical relevance in the anatomical scan (n=1). The final sample of 15 included 9 females / 6 males, aged 18 – 34 years (*M* = 21.6 years; *SD* = 4.1 years). We note that this final sample was collected after study preregistration and therefore does not include the three pilot participants upon whose data the preregistration was based. The Human Subjects Institutional Review Board of the University of Wisconsin–Madison approved the study protocol, and all participants provided informed consent.

### Visual stimuli and behavioral tasks

#### Load-of-3 trials

The stimuli consisted of sinusoidal gratings. The contrast of the gratings was held constant at 0.6 and the spatial frequency was 1 cycle/deg, phase angle varying randomly between 0 deg and 179 deg for each presentation. The gratings were presented within a circular patch with a 4 deg diameter, at one of six locations around a central fixation circle (156 pixels in diameter). The six stimulus locations were positioned at seven degrees eccentricity from fixation at angles of 0 deg, 60 deg, 120 deg, 180 deg, 240 deg, and 300 deg (see Figure 1A for a schematic of the stimulus configuration). For each presentation, the gratings were randomly assigned one of six orientation values: 10 deg, 40 deg, 70 deg, 100 deg, 130 deg, or 160 deg, with a random jitter between −3 deg and + 3 deg. The cardinal orientations were not included in the set to reduce the likelihood of verbal encoding. Probe stimuli consisted of a grating with a superimposed digit centered in the patch (40 pixels in height), rendered in red. On 50% of trials (‘match’), probe stimuli were presented with the same orientation as the trial’s randomly chosen target grating; on the other 50% of trials (‘non-match’), the orientation of the probe stimulus was defined as the original sample orientation (i.e., the ‘match’ orientation) jittered by a randomly selected amount from the set of [−25, −20, −15, −10, 10, 15, 20, 25] deg. Therefore, non-match orientations never overlapped with the other sample orientations (but could be within 2 deg). All stimuli were generated and presented in MATLAB (MathWorks) and Psychtoolbox-3 (http://psychtoolbox.org; Brainard 1997; Pelli 1997; Kleiner et al. 2007).

**Figure 1.**
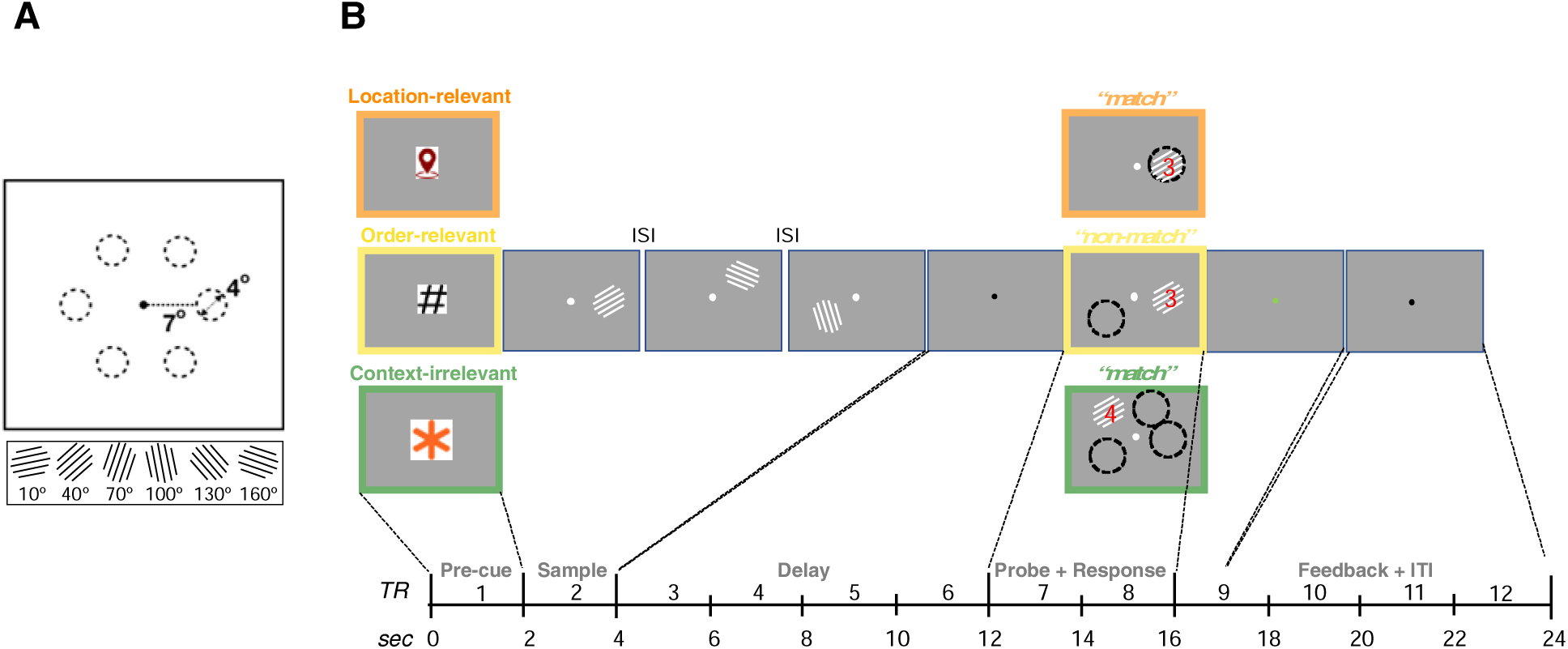
Experimental paradigm. (**A)** Sample stimuli consisted of oriented-grating patches (4 deg diameter) drawn from a fixed set of six values and presented at three from a fixed set of six equally-spaced locations, each at 7 deg eccentricity relative to a central fixation point. **(B)** The primary task comprised three interleaved trial types that each began with a pre-cue indicating trial type. The pre-cue was followed by the presentation of three sample gratings of different orientations, presented serially, each at a different location. After an 8-second delay period, the probe stimulus appeared. The basis for the recognition decision was determined by the trial type: on location-relevant trials, subjects compared the orientation of the probe to their memory of the orientation of the sample item that had appeared at the location occupied by the probe; on order-relevant trials, subjects compared the orientation of the probe to their memory of the orientation of the sample item that occurred in the ordinal position indicated by the superimposed digit; on context-irrelevant trials, subjects compared the orientation of the probe to their memory of all three sample item orientations. For illustration purposes, this figure uses dashed circles to illustrate the location of the sample(s) being tested; these did not appear during the experiment.

Each trial of the delayed-recognition task began with an instructional cue that identified the trial type – *location-relevant* (a “location pin” icon), *order-relevant* (“#”), or *context-irrelevant* (“*”) – followed by the serial presentation of three sample stimuli (500 ms presentation, 250 ms inter-stimulus interval (ISI)), each appearing at a different location. After an 8-second delay, a probe stimulus (oriented grating with same properties as the sample stimuli, but with a superimposed digit rendered in red) appeared for 4 sec, and a “match” or “nonmatch” response was required while the probe remained on the screen. Feedback (green fixation dot = correct; red fixation = incorrect or time-out) was provided for the first sec following probe offset, after which the fixation dot was black for the remaining 7-sec of the ITI (Figure 1B). On *location-relevant* and *order-relevant* trials, the probe appeared in one of the three locations where a sample had appeared, and the superimposed digit was “1”, “2”, or “3”. On *location-relevant* trials, the probe’s location was the same as the sample against which it should be compared, whereas on *order-relevant* trials the digit indicated the ordinal position of the sample (i.e., the one that appeared first, second, or third) against which it should be compared. On these two trial types, the value of the irrelevant dimension was selected at random from the remaining samples, meaning that the probe’s location and digit never cued the same sample. On *context-irrelevant* trials the probe stimulus appeared at one of the three locations that had not been occupied by a sample and the superimposed digit was randomly chosen from the set of “4”, “5”, and “6” (i.e., neither corresponded to any of the samples), and the subject was to indicate if the probe orientation matched that of any of the three samples (Figure 1B). For each trial type, the orientation of the probe matched the orientation of the critical sample(s) on 50% of the trials. Subjects responded via button press on the keyboard (“1” for match, “2” for non-match).

#### Load-of-1 trials

The stimuli and procedure for the load-of-1 trials were similar to those of the load-of-3 trials, with three exceptions: No precue was presented; the 500 ms presentation of the single sample item was followed by an 8.5-second delay, and the recognition probe consisted solely of a grating (i.e., no digit) and always appeared at the same location as had the sample.

### Experimental procedures

#### Behavioral training/screening session

The behavioral training/screening session took place on a separate day prior to fMRI scanning. It began with a block of 18 load-of-1 trials to familiarize subjects with the delayed recognition task. The session continued with six blocks of 18 load-of-3 trials. Subjects achieving an accuracy of 83% or above in each of the three context conditions (i.e., *location-relevant*, *order-relevant*, *context-irrelevant*) in at least one of the load-of-3 blocks were invited to participate in the fMRI sessions, which took place on the soonest dates afforded by the subject’s and scanner availability. Enrollment continued until the planned sample of 15 subjects completed both fMRI sessions and the data for the sample were all deemed useable (i.e., they were free of significant artifacts and other image quality concerns).

#### fMRI sessions

The behavioral tasks completed during fMRI scanning were identical to those the subject completed during the behavioral training/screening session. The experimental stimuli were presented using a 60-Hz projector (Silent Vision 6011; Avotec) and viewed through a coil-mounted mirror. The viewing distance was 68.58 cm and the screen width was 33.02 cm. fMRI scanning occurred in two sessions per subject. The first fMRI session consisted of eight 18-trial runs of load-of-3 trials followed by five 18-trial runs of load-of-1 trials. Each run lasted for 7 minutes and 20 seconds. The second fMRI session consisted of seven 18-trial runs of Load-of-3 trials followed by five 18-trial runs of load-of-1 trials. The two sessions combined yielded a total of 270 load-of-3 trials (90 per context-binding trial type; 15 per location per context-binding trial type) and a total of 180 load-of-1 trials (30 per location). Responses were given via button press using two buttons on an MR-compatible 4-button box. The buttons corresponding to a ‘match’ or ‘non-match’ response were swapped and counterbalanced between subjects.

Whole brain images were acquired with a 3-T MRI scanner (Discovery MR750; GE Healthcare) at the Lane Neuroimaging Laboratory at the University of Wisconsin–Madison. For all subjects, a high-resolution T1-weighted image was acquired with a fast-spoiled gradient-recalled echo sequence [repetition time (TR) 8.2 ms, echo time (TE) 3.2 ms, flip angle 12°, 172 axial slices, 256 x 256 in-plane, 1.0 mm isotropic]. A T2*-weighted gradient echo pulse sequence was used to acquire data sensitive to the BOLD signal while subjects performed the delayed recognition task (TR 2,000 ms, TE 25 ms, flip angle 60°, within a 64 x 64 matrix, 40 sagittal slices, 3.5 mm isotropic). Each of the 25 fMRI scans generated 220 volumes. Eye movements were monitored using a ViewPoint EyeTracker system (Arrington Research).

fMRI data were preprocessed using the Analysis of Functional Neuroimages (AFNI) software package (https://afni.nimh.nih.gov; Cox 1996). Each run began with eight seconds of dummy pulses to achieve a steady state of tissue magnetization before task onset. All volumes were spatially-aligned to the first volume of the first run using a rigid-body realignment and then aligned to the T1 volume. Volumes were corrected for slice-time acquisition, and linear, quadratic, and cubic trends were removed from each run to reduce the influence of scanner drift. For univariate analyses, data were spatially smoothed with a 4-mm full-width at half-maximum Gaussian and z-scored separately within run for each voxel. For multi-voxel pattern analyses (MVPA) and inverted encoding modeling (IEM), data were z-scored separately within runs for each voxel, but the data were not smoothed. All analyses were carried out in each subject’s native space.

Univariate analyses entailed calculating the percentage signal change in BOLD activity relative to baseline for each time point during the delayed-recognition task. The average BOLD activity of the first TR of each trial was used as baseline. A conventional mass-univariate general linear model (GLM) analysis was implemented in AFNI, with sample, delay, and probe periods of the task modeled with boxcars (2 s, 8 s, and 4 s in length, respectively) convolved with a canonical hemodynamic response function (HRF). Differences in BOLD activity from baseline were evaluated with one-sample *t*-tests and Bayes factors of the likelihood of the different-from-baseline alternative vs. not-different-from-baseline null hypothesis.

To generate regions of interest (ROIs), we followed the approach used by Gosseries, Yu, et al. (2018) and Cai et al. (2019), and focused our analyses on two anatomically constrained functional ROIs: an *occipital-sample* ROI and a *parietal-delay* ROI. The *occipital-sample* ROI was defined as the 500 voxels displaying the strongest loading on the contrast [sample – baseline] from the GLM, collapsed across the three context-binding conditions in the load-of-3 trials, and located within the anatomical mask for occipital cortex from the Talairach Daemon atlas for AFNI transformed to each subject’s individual structural image via affine transformations (Jenkinson & Smith 2001), and further refined via nonlinear interpolation (Andersson et al. 2007). The *parietal-delay* ROI was defined as the 500 voxels displaying the strongest loading on the contrast [delay – baseline], also collapsed across the three context-binding conditions and located within an anatomical mask for parietal cortex from the Talairach Daemon atlas for AFNI transformed to each subject’s individual structural image using the same procedures as used for defining the standard occipital mask.

In addition to the ROI generation described above, we also defined anatomical ROIs for subregions of the intraparietal sulcus (IPS0-IPS5) based on the Wang et al. (2015) probabilistic atlas, and selected the 500 most responsive voxels within each (see Gosseries, Yu, et al. 2018). This approach allowed more granularity in some of the tests of the activity related to context-binding demands in IPS.

### Data analysis

#### Analysis of behavioral data

Behavioral performance during the fMRI portion of the study was analyzed for accuracy and reaction time. Accuracy was quantified as percentage of trials in which a correct response (‘match’ or ‘non-match’) was given. Accuracy in each of the four trial types was tested against the chance performance level of 50% using one-tailed, one-sample *t*-tests. Two-tailed paired-sample *t*-tests were used to test for differences between conditions. Both within-subject and between-subject reaction times were assessed as median reaction times due to positive skew in the distributions. 95% confidence intervals for the median between-subject reaction times were obtained with a bias-corrected and accelerated (BCa) bootstrapping procedure with 10,000 iterations, implemented using the *BCa_bootstrap.m* MATLAB function (Van Snellenberg 2018). Two-sided sign tests using the exact method for obtaining the *p*-value were used to test for differences in median reaction times between conditions.

#### Implementation of hypothesis tests

##### Context binding versus load (Hypothesis 1)

These analyses were motivated by the idea that delay-period load sensitivity of activity in posterior parietal cortex, including IPS, may reflect, at least in part, the demands on context binding that often covary with memory load (Gosseries, Yu, et al. 2018; Cai et al. 2019; Cai et al. 2020). Its tests were implemented with univariate analyses that tested for task vs. baseline and context-relevant vs. *context-irrelevant* differences in late-delay period BOLD (TRs 6-7), via one-sample *t*-tests and Bayes factors of the likelihood of the conditional differences (alternative) hypothesis vs. no differences (null) hypothesis.

In addition to the a priori hypothesis tests described above, we also planned several additional analyses that, although not directly testing the three sets of hypotheses that were the primary motivation for this work, could nonetheless provide further insight into the role of context-binding in VWM. Related to the question of dissociating sensitivity to context-binding versus to load, we also planned to carry out whole-brain contrasts of delay-period activity for context-relevant trials versus *context-irrelevant* trials and for *location-relevant* versus *order-relevant* trials. The delay period was modeled in a GLM with 8s boxcar regressors spanning the delay period and convolved with a canonical hemodynamic response function (coded by trial type), and six nuisance regressors related to movement-related artifacts; sample and probe events were not included in the model. Parameter estimates for the delay-period regressors were calculated from the least mean squares fit of the model to the data. To test for statistical significance of differences between conditions, we first formed subject-specific contrast, then normalized the resulting contrast images to a standard Montreal Neurological Institute (MNI) space, then submitted the set of 15 images (one per subject) to AFNI’s 3dttest++ with the input “-ClustSim”, which carried out a permutation analysis to compute a cluster-size threshold for a given voxel-wise *p*-value threshold, such that the probability of any clusters surviving the dual thresholds is at some given level. Results are reported after applying a threshold of *p* < 0.001 uncorrected in conjunction with a prescribed cluster size of 39 voxels (3.5 mm) for the context-relevant vs. *context-irrelevant* analysis and 40 voxels for the *location-relevant* vs. *order-relevant* analysis to achieve *p* < 0.05 familywise error correction for multiple comparisons across the whole brain volume. (Due to an oversight, the preregistered analysis plan did not explicitly state that we also planned to carry out these analyses in IPS subregions. However, because the Methods section of the preregistered document does indicate that we would analyze the activity in the IPS subregions, and to improve narrative flow, we will report the results of these analyses from the IPS subregions immediately after the whole-brain results.)

##### Domain specificity of context binding (Hypothesis 2)

These analyses assessed evidence that patterns of activity in brain areas associated with VWM would be sensitive to (i) the level of demand on context binding, and (ii) the informational domain of task-specific context. To test this, we carried out trialwise category-level multivariate voxel pattern analysis (MVPA) to discriminate activity observed on (i) context-relevant vs. *context-irrelevant* trials; and (ii) *location-relevant* vs. *order-relevant* trials in occipital and parietal ROIs. MVPA was performed via L2-regularized logistic regression with a penalty term of 25, using the Princeton Multi-Voxel Pattern Analysis toolbox (www.pni.princeton.edu/mvpa/). The classifiers were trained and tested on the patterns corresponding to the two categories (i.e., context-relevant vs. *context-irrelevant* and *location-relevant* vs. *order-relevant*) at each time point (TR) through a leave-one-trial-out k-fold cross-validation procedure for each subject and ROI separately. We carried out two versions of the context-relevant vs. *context-irrelevant* MVPA. First, because there were twice as many context-relevant trials as *context-irrelevant* trials, we randomly selected half the *location-relevant* and half the *order-relevant* trials and trained a classifier to discriminate these context-relevant trials from *context-irrelevant* trials, then repeated this process 100 times and averaged performance across iterations to obtain a measure of classifier accuracy within the given ROI for each subject. In the second approach, one classifier was trained to decode *location-relevant* trials from *context-irrelevant* trials, and a second classifier was trained to decode *order-relevant* trials from *context-irrelevant* trials. This approach allowed us to use all trials, summarizing classifier performance using the average performance of the two classifiers. Because the two approaches yielded nearly identical results, we report the results of the latter.

Classifier performance was summarized using the area under the curve (AUC). AUC reflects the sensitivity of the classifier in discriminating between the two categories and was computed as follows: We first selected a target category for each of the two classifiers and computed the proportion of hits vs. false alarms. The AUC was then computed using trapezoidal approximation to estimate the area based on these two proportions. An AUC greater than 0.5 indicates sensitivity to the target category. Statistical results were summarized through one-tailed one-sample *t*-tests comparing the classifier AUC against 0.5. False discovery rate (FDR) correction was used to correct the *p*-values for the 12 comparisons against chance level (i.e., at each TR) for a given ROI and classifier.

In addition to the a priori hypothesis tests described above, we also planned to carry out whole-brain searchlight MVPA comparisons of context-relevant vs. *context-irrelevant* and *location-relevant* vs. *order-relevant* trial types. For these analyses, we used The Decoding Toolbox (TDT; Hebart et al. 2015) submitting the beta images resulting from GLMs similar to *Additional, Hyp 1*, with the exception that the models were re-fit to each run separately and spanned the 14-s time frame from sample through probe, resulting in a set of betas for each condition and run. We ran cross-validated leave-one-run-out searchlight decoding analyses, wherein a separate support vector machine was built for each voxel, fitted to the beta values within a sphere with a radius of 3 voxels. This resulted in three-dimensional decoding accuracy maps in native space for each participant and analysis. (Decoding accuracy was calculated relative to chance level (i.e., 50% was subtracted from all accuracies)). These individual-subject maps were normalized into MNI space and, to identify significant classification performance, the set of 15 images (one from each subject) was submitted to AFNI’s 3dttest++ with the input “ClustSim”. Results are reported after applying a threshold of p < 0.001 uncorrected in conjunction with a prescribed cluster size of 16 voxels (3.5 mm) for the context-relevant vs. *context-irrelevant* contrast and 14 voxels for the *location-relevant* vs. *order-relevant* contrast to achieve p < 0.05 familywise error correction for multiple comparisons across the whole brain volume.

##### Controllability of context binding (Hypothesis 3)

These analyses operationalized the idea that the representation of stimulus context in VWM is under strategic control by positing differences in the strength of inverted encoding model (IEM) reconstruction of the neural representation of the location of the three sample items as a function of their relevance for retrieval from VWM. To test this, we used multivariate IEMs (Brouwer & Heeger 2009, Brouwer & Heeger 2011; Serences & Saproo 2012; Sprague & Serences 2013; Sprague et al., 2018; Sprague et al., 2019) to reconstruct channel tuning function (CTF) response profiles that track the perceived and remembered probe locations from multivoxel patterns of activity. The choice of the IEM approach was motivated by the fact that it allowed us to obtain an estimate of the strength of the representation of the location of each of the three sample items in working memory during the delay and at probe (Sprague et al., 2019; for further discussion of IEM model assumptions and best practices, see also Sprague et al., 2018a, 2019; Adam & Serences, 2021).

IEMs were trained on sample-evoked signal (TR 4) from load-of-1 trials, labeled by location, and tested on load-of-3 trials. fMRI data from all trials (both correct and incorrect) were included in the IEM training and reconstruction. We extracted the normalized responses of each voxel in the *occipital-sample* ROI for each time point after *z*-scoring within each run. The logic behind the IEM method is that BOLD signal from each voxel can be construed as a weighted sum of responses from six hypothetical channels optimally tuned for a specific stimulus location (i.e., 0 deg, 60 deg, 120 deg, 180 deg, 240 deg, and 300 deg). For each IEM, we estimated the weight matrix (*W*) that projects the hypothesized channel responses (*C_1,_*, *k* x *n; n*: the number of repeated measurements; *k:* the number of locations) to the actual measured fMRI signals in the load-of-1 training data set (*B*_1_, *v* x *n*, v: the 500 voxels in the *occipital sample* ROI). This relationship was characterized by:

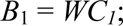

where *W* was the weight matrix (*v* x *k*).

The least-squares estimate of the weight matrix (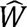) was calculated using linear regression:

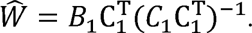

We then inverted this weight matrix to estimate channel responses (*Ĉ*_2_) for test data from each load-of-3 test trial (*B_2_*):

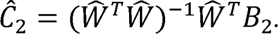

The average response output for each channel across trials was obtained by circularly shifting each response to a common center of 0°. The shifted channel outputs were averaged across all iterations in each subject. To quantify the resultant reconstructions of neural representations of stimulus location, we collapsed over channel responses on either side of the target channel (i.e., channel = 0° after shifting the outputs), averaged, and then used linear regression to estimate slope of the reconstruction for each subject at each tested TR. Finally, we computed the between-subjects average slope. A slope value greater than 0 can be interpreted as evidence for an active neural representation (Foster et al. 2017), and in these analyses slope served as a proxy for the strength of the representation. Statistical significance of the slope was assessed with a bootstrapping method (Ester et al. 2015, Ester et al. 2016). We randomly selected (with replacement) a set of reconstructions equal to the sample size and averaged them. This step was repeated 2500 times. We estimated the slope of each reconstruction, and a *p-*value was computed as the proportion of permutations for which the slope estimates were ≤ 0 for positive reconstructions. For negative reconstructions, the *p*-value was computed as 1 – the proportion of permutations for which the slope estimates were ≤ 0. Additionally, a 95% confidence interval for slope was constructed using the lower 2.5^th^ and upper 97.5^th^ percentiles of the bootstrapped distribution. Comparison of reconstruction slopes across trial types -- critical for the tests of our hypotheses – was carried out with two-sided paired-sample t-tests and Bayes factors. (Note that, because the neural coding of the representation of ordinal position is poorly understood (relative to egocentric location), this approach was limited to studying the controllability of location context.)

To test the sub-hypotheses under *Hypothesis 3*, the IEM trained on the sample-evoked signal from the load-of-1 trials was used to reconstruct: (*Hyp. 3A*) the location of the probe (TRs 9 and 10); (*Hyp. 3B*) the location of the sample cued by the superimposed digit in the probe stimulus from TRs 9 and 10, (*Hyp. 3C*) the location of the sample that corresponded to neither the location of the probe nor the sample location cued by the superimposed digit, from TRs 9 and 10. Hyp. 3A operationalizes the assessment of the sensitivity of the processing of the physical properties of the probe (specifically, the encoding of its location) to strategy (i.e., the probe’s location is not relevant on *order-relevant* and *context-irrelevant* trials). *Hyp. 3B* operationalizes a test of whether an item’s context is reinstated when that item is cued for the recognition judgment. *Hyp. 3C* operationalizes a test of whether the context of all items in the memory set are reinstated nonspecifically (an outcome that would argue against the strategic control of stimulus context in VWM). Note that, because the results generally did not differ between TRs 9 and 10, we averaged the results for each subject to obtain a single reconstruction slope for visualization purposes.

In addition to the hypothesis-testing analyses described above, we also planned three additional analyses. (*i*) The first entailed repeating the analyses testing *Hypothesis 3* but with IEMs trained on the probe-evoked signal (TRs 9 and 10) of load-of-3 trials, and labeled according to the location-on-the-screen of the probe. (*ii*) The second entailed using IEMs trained on the sample location-evoked signal (TR 4) from load-of-1 trials from the *occipital-sample* and *parietal-delay* ROIs, and tested on late delay-period signal (TR 6) within the same ROI from load-of-3 trials. (*iii*) The third entailed IEM of stimulus orientation (i.e., not location). IEMs were trained on sample-evoked signal (TR 4) from load-of-1 trials, labeled by sample orientation, and tested on each time point (TR) of the load-of-3 trials, labeled according to the to-be probed sample orientation.

##### Post hoc analyses (exploring the controllability of context binding (Hypothesis 3))

The preregistered analyses designed to assess the controllability of context binding, described in the previous subsection, were based on the assumption that IEMs trained on sample-evoked signal (at TR 4) from load-of-1 trials would be able to reconstruct the representation of the location of items other than the probe during the epoch when the probe was on the screen. However, as will be seen below, these analyses failed to produce interpretable results (and thus constituted a failure of perception-based models to generalize to VWM). These outcomes prompted us to carry out two post hoc analyses in which we modified the IEM procedure by training and testing IEMs on the same time points in the trial. These post hoc analyses did yield informative results. For *post hoc analysis 1*, the analyses comprised a modification of the late-delay period analysis (ii) described in the previous paragraph. It involved training an IEM on the late-delay period signal (TR 6) from load-of-1 trials from the occipital-sample and parietal-delay ROI, labeled by the sample location and testing on the signal from the load-of-3 trial types at the same TR. For *post hoc analysis 2,* the analyses comprised a modification of the probe-period analysis (i) described in the previous paragraph. (i.e., leave-one-run-out cross-validation in which we trained and tested on the load-of-3 trial TRs 9 and 10). Whereas for probe-period analysis (i) the training data were labeled according to the location-on-the-screen of the probe, for *post hoc analysis 2* we trained four IEMs, labeling the training data differently for each one: IEM_post hoc 2 #1_ – trained and tested on the probe’s location on the screen; IEM_post hoc 2 #2_ – trained and tested on the location of the item referenced by the superimposed digit in the probe stimulus (note that this analysis is undefined for *context-irrelevant* trials, in which the superimposed digit ranged from 4-6); IEM_post hoc 2 #3_ – trained and tested on the location of the item that was not cued by the probe stimulus (note that, for *context-irrelevant* trials, this includes all three sample locations); IEM_post hoc 2 #4_ – trained and tested on the locations not occupied by a sample (there were three of these on every trial). These analyses were carried out in both the *occipital-sample* and *parietal-delay* ROIs. In the event that either IEM_post hoc 2 #2_ or IEM_post hoc 2 #3_ was successful, IEM_post hoc 2 #4_ would serve as a control to confirm that this approach of training and testing would not be able to reconstruct, at TRs 9 and 10, the locations that had not been occupied by a sample item.

## Results

### Behavior

Recognition accuracy during fMRI scanning was significantly higher on load-of-1 than on load-of-3 trials (*t*(14) = 15.121, *p* < .001) and median reaction time was significantly faster on load-of-1 than on load-of-3 trials (*exact binomial p =* 0.0074; two-tailed sign rank test). Within load-of-3 trial types, accuracy was significantly greater for *location-relevant* than for both *order-relevant* (*t*(14) = 5.019, *p* < .001) and *context-irrelevant* (*t*(14) = 4.797, *p* < .001) trials. Note that although accuracy on *context-irrelevant* trials did not differ statistically from chance (*t*(14) = 1.470, *p* = .082), it also did not differ statistically from accuracy on *order-relevant* trials (*t*(14) = 0.899, *p* = .384). At the level of individual subjects, binomial tests indicated that the accuracy on *context-irrelevant* trials was greater than chance for 7 of the 15. This prompted us to re-examine behavior in the three pilot subjects on whom our preregistered hypotheses were based, and binomial tests indicated that the accuracy on context-irrelevant trials was greater than chance for 2 of the 3.

Additionally, within load-of-3 trial types, median reaction time was significantly slower for *order-relevant* trials than both *location-relevant* (*exact binomial p =* 0.0352) and *context-irrelevant* (*exact binomial p =* 0.0074) trials, and did not differ between *location-relevant* and *context-irrelevant* trials (*exact binomial p =* 1; Figure 2).

**Figure 2.**
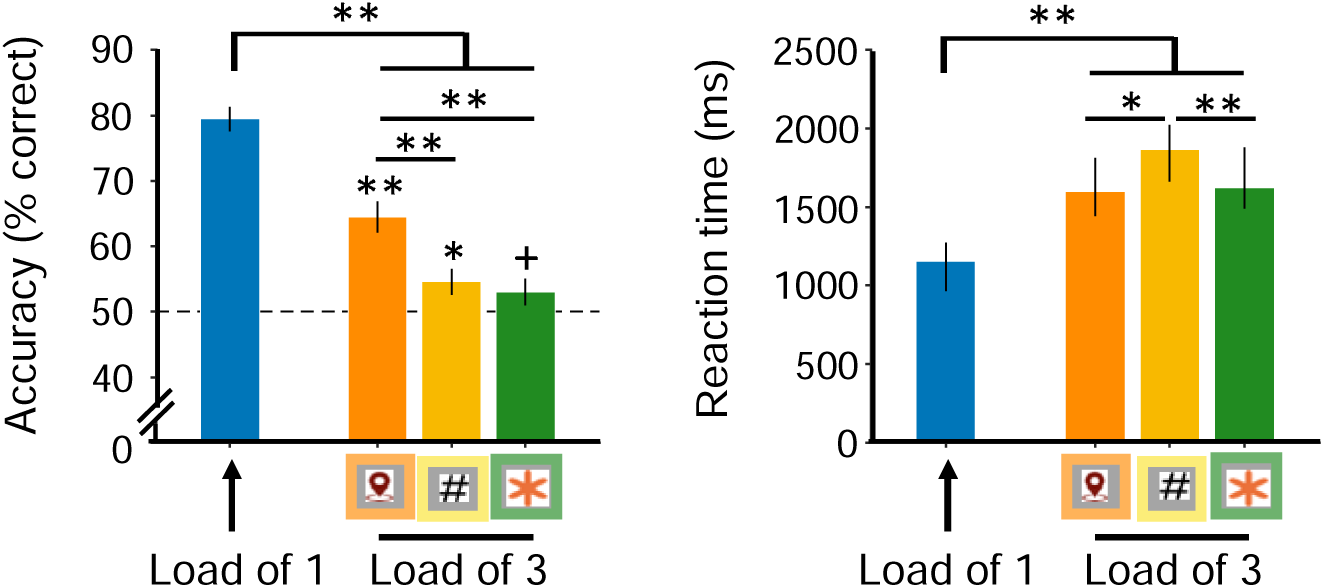
Behavioral results. Left panel: Accuracy as a function of trial type. Error bars correspond to +/−1 standard error of the mean (SEM). Chance performance = 50% correct. Right panel: Reaction time as a function of trial type. Error bars correspond to bootstrapped 95% confidence intervals for the median. ** p < .01; * p < .05; + p < .1.

### fMRI

Context binding vs. load *(Hypothesis 1)*. Focusing first on PPC *(Hypotheses 1A and 1C)*, univariate analyses of BOLD signal activity in the *parietal-delay* ROI confirmed the expected elevation of activity at the late delay-period TRs (6-7) for all trial types (all *t-statistics* ≥ *3.943,* all *p-values* < .002, all *BF_10_* ≥ 27.806; Figure 3A). Late delay-period activity was modulated by context-binding demands, with greater activity for context-relevant trials than *context-irrelevant* trials (*t*(14) = 1.928, *p* = .037, *BF_10_* = 2.187) and load-of-1 trials (*t*(14) = 3.717, *p* < .001, *BF_10_* = 48749). To address the possibility that this difference may have been driven by the subjects with chance-level performance on context-irrelevant trials, we repeated this analysis with only the 7 subjects whose performance on these trials was above chance, and observed that although late delay-period activity was numerically greater for context-relevant than context-irrelevant trials, this difference no longer achieved threshold for significance (t(6) = 1.288, *p* = .123, *BF_10_* = 1.148). Finally, to partly offset the low sample size, we carried out this analysis a third time, after adding in data from the two pilot subjects whose performance on context-relevant trials exceeded chance, and the results exceeded the threshold for significance (t(8) = 2.131, *p* = .033, *BF_10_* = 2.879). Additionally, inspection of data from individual subjects revealed two subjects whose delay-period signal was an average of 0.21% greater for *context-irrelevant* than context-relevant trials. This is in comparison with the remaining subjects for whom this signal was an average of 0.13% greater on context-relevant than *context-irrelevant* trials.

**Figure 3.**
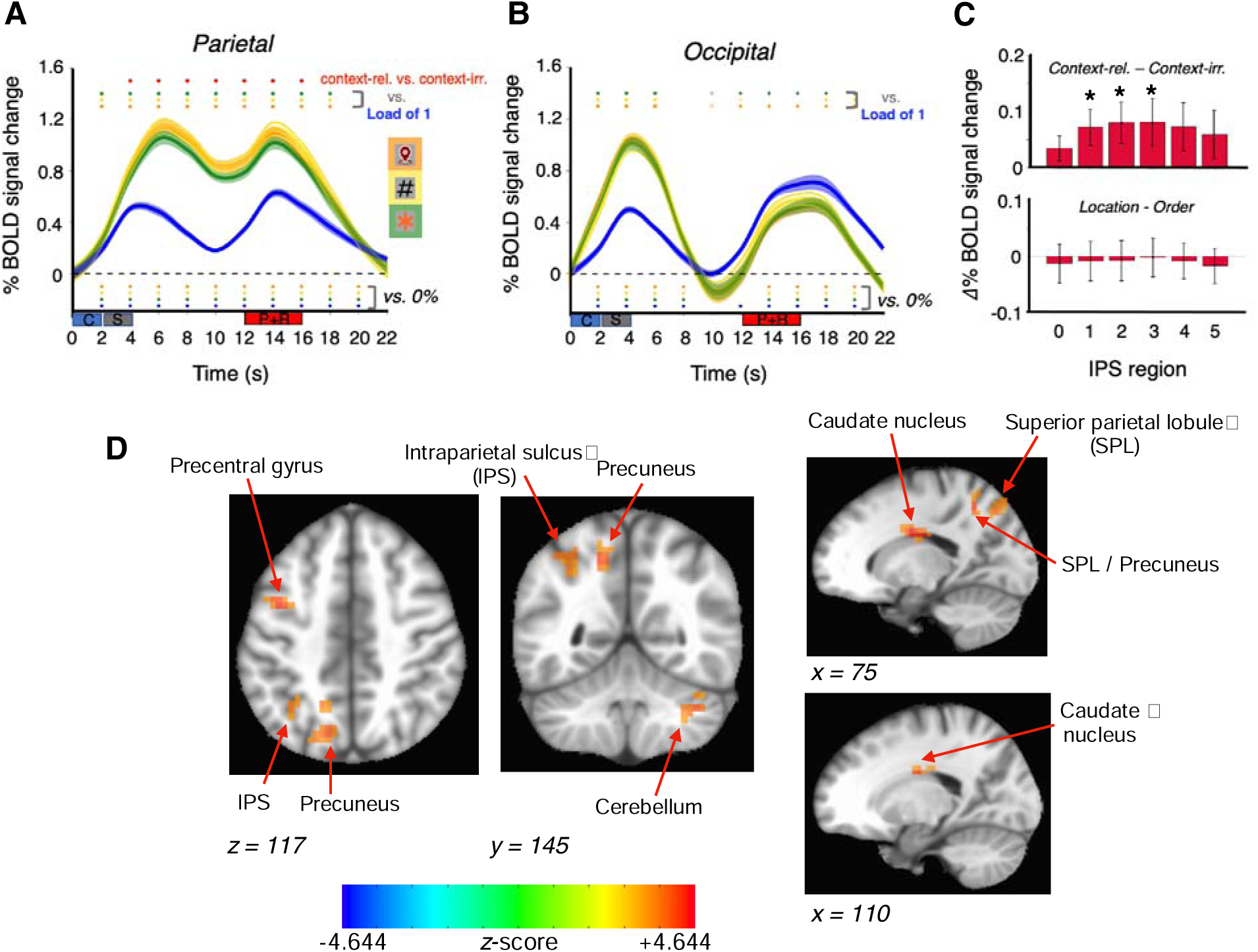
Univariate analysis results. **(A)** Percent BOLD signal change in the *parietal-delay* ROI at each time point for the load-of-1 (blue), *context-irrelevant* (green), *order-relevant* (yellow), and *location-relevant* (orange) trial types. Dark lines correspond to the mean at each time point; translucent ribbons correspond to +/−1 SEM (time courses smoothed for visualization). Symbols below the dashed line indicate significance relative to baseline; symbols above the data indicate significant differences between conditions (note that *Hypothesis 1* relates to TRs 6-7; 10-14 s). Rectangles below the *x*-axis denote the cue (‘C’), sample (‘S’) and probe + response (P + R) events. **(B)** Same format as **(A)** for the *occipital-sample* ROI. **(C)** Difference in BOLD signal change between the (Top) context-relevant and *context-irrelevant* trials types and (Bottom) *location-relevant* and *order-relevant* trial types during the late delay period (TRs 6-7; 10-14 s) for individual subregions of IPS. Error bars correspond to +/−1 SEM. * *p* < .05. **(D)** Clusters identified in a whole-brain analysis contrasting delay-period activity for context-relevant vs. *context-irrelevant* trial types (see also Supplementary Figure 1). Positive z-scores correspond to context-relevant > *context-irrelevant*.

This pattern was observed in three subregions of IPS, with significantly greater late-delay period activity for context-relevant trials than *context-irrelevant* trials in IPS 1-3 (all *t-statistics* ≥ 1.9039, all *p-values* ≤ .0388, all *BF_10_* ≥ 2.1126; remaining subregions all *t-statistics* ≤ 1.7105, all *p-values* ≥ .0546, all *BF_10_* ≤ 1.6150; Figure 3C). These results were consistent with *Hypothesis 1A*. Furthermore, in the PPC no differences were identified in delay-period activity when the two context-relevant trial types were compared, either in *parietal-delay* ROI (all *t-statistics* ≤ 0.4311, all *p-values* ≥ .4659, all *BF_10_* ≤ 0.2747), or in any subregion of IPS (all *t-statistics* ≤ 0.5453, all *p-values* ≥ .5941, all *BF_10_* ≤ 0.2990; Figure 3C; these results were consistent with *Hypothesis 1C*).

Turning next to occipital cortex (*Hypotheses 1B and 1C)*, in the *occipital-sample* ROI delay-period BOLD activity did not differ from baseline for all trial types (all *t-statistics* ≤ 1.47, all *p-values* ≥ .1637, all *BF_10_* ≤ 0.6406), and activity in all three of the load-of-3 trial types was greater than the activity in the load-of-1 trial type during the very early delay period (TR 4; 6-8 s; all *t-statistics* ≥ 6.965, all *p-values* < .001, all *BF_10_* ≥ 3201.7; Figure 3B) but not at later time points. These results were consistent with *Hypothesis 1B*. Furthermore, in the *occipital-sample* ROI no differences were identified in delay-period activity when the two context-relevant trial types were compared (all *t-statistics* ≤ 0.465, all *p-values* ≥ .6755, all *BF_10_* ≤ 0.1873; consistent with *Hypothesis 1C*).

For the whole-brain analysis of delay-period activity, the context-relevant vs. *context-irrelevant* contrast revealed six clusters showing context-relevant > *context-irrelevant*– left precuneus, left superior parietal lobule, left precentral gyrus, right cerebellum, and caudate nucleus bilaterally (see Figure 3D; **Supplementary Figure 1**; Table 2). For the *location-relevant* vs. *order-relevant* contrast, no clusters survived thresholding.

**Table 2.**
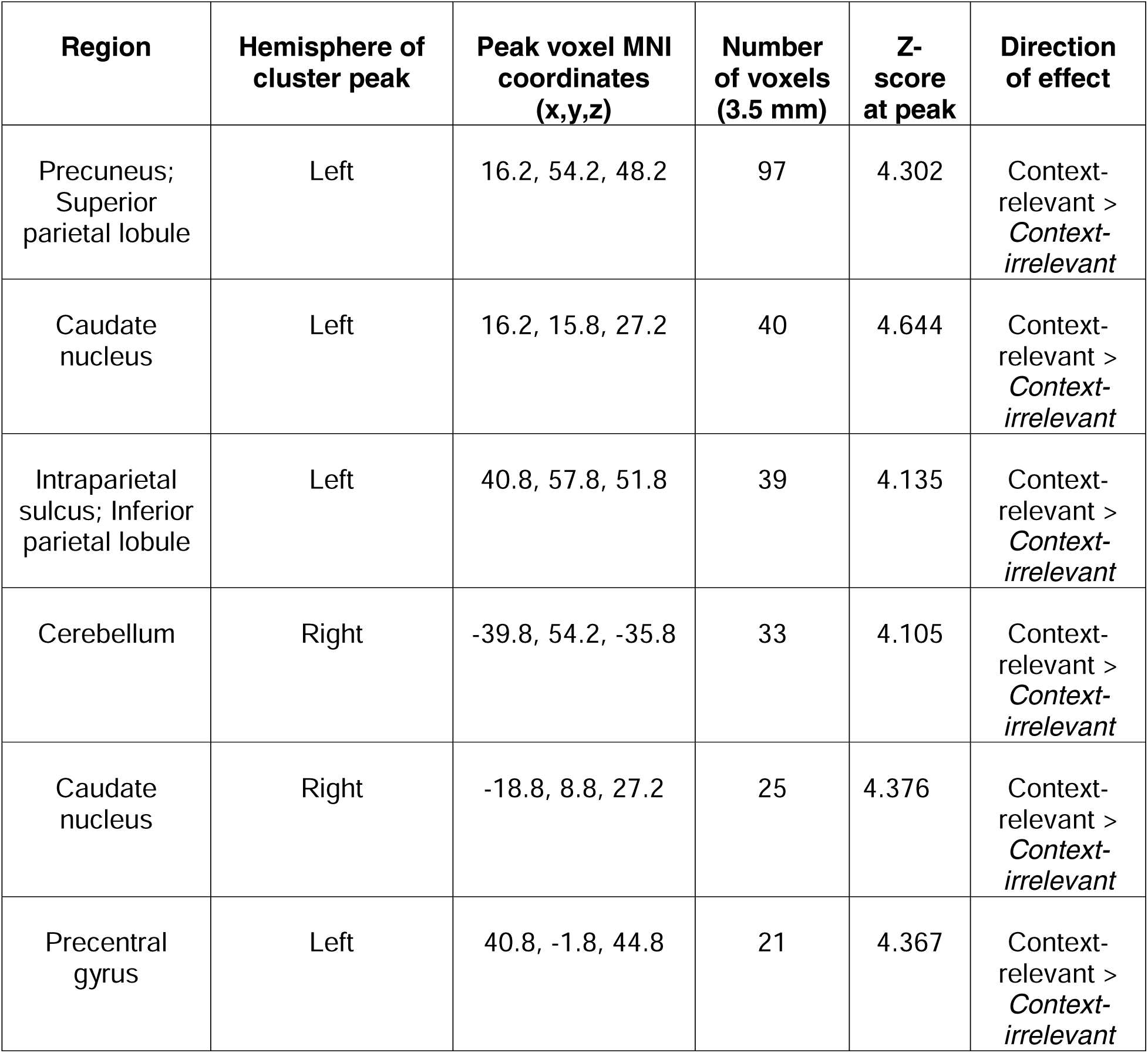
Regions identified in whole-brain contrast of delay-period activity for context-relevant vs. *context-irrelevant* trial types (see Figure 3.D and Supplementary Figure 1).

Domain specificity of context binding *(Hypothesis 2)*. In the *parietal-delay* ROI, MVPA successfully classified context-relevant from *context-irrelevant* trials, building steadily from TR3 (4-6 s) through the remainder the trial. In the *occipital-sample* ROI, the temporal profile of classifier performance was reversed – strongest early in the trial, then declining to chance levels during the late delay period (TR 7; 12-14 s) and fluctuating thereafter; Figure 4A). Classification of *location-relevant* from *order-relevant* trials in the *parietal-delay* and *occipital-sample* ROIs followed qualitatively similar patterns (Figure 4B). These results were consistent with *Hypothesis 2*.

**Figure 4.**
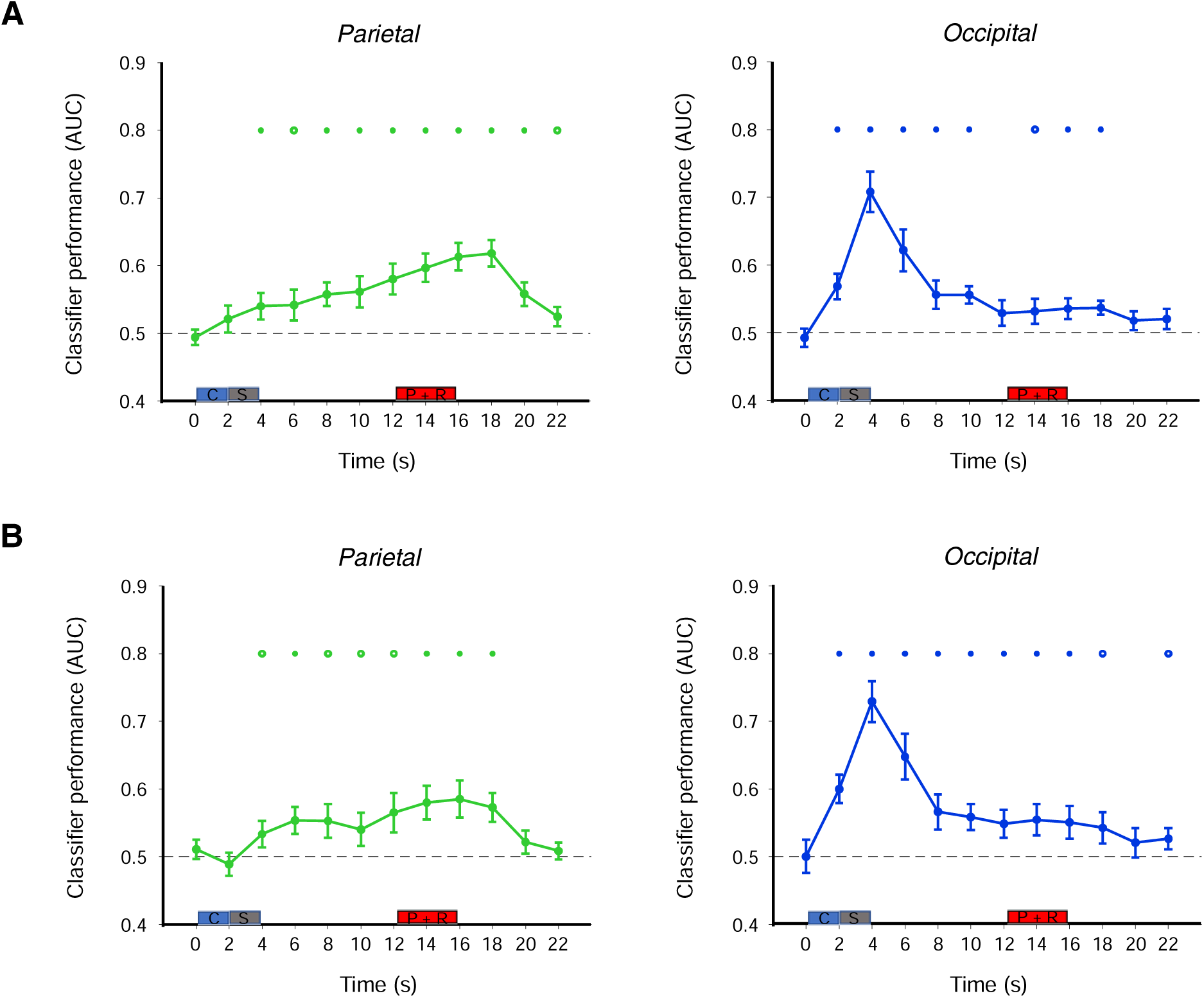
MVPA results. **(A)** Time course of classifier accuracy (AUC) in discriminating context-relevant from *context-irrelevant* load-of-3 trial types in the *parietal-delay* ROI (left) and *occipital-sample* ROI (right). **(B)** Time course of classifier accuracy (AUC) in discriminating *location-relevant* from *order-relevant* load-of-3 trials in the *parietal-delay* ROI (left) and *occipital-sample* ROI (right). Rectangles denote the pretrial instructional cue (‘C’), sample (‘S’) and probe + response (‘P + R’) events. Error bars correspond to +/−1 SEM. Circular symbols indicate significance relative to chance level performance (0.5); solid symbols: fdr-corrected *p*-values < .05; open symbols: fdr-corrected *p*-values < .1.

Next, to look more broadly at the representation of stimulus context across the brain, we carried out a searchlight analysis, collapsing across the entirety of the trial. Results indicated that context-relevant trials could be discriminated from *context-irrelevant* trials in areas that overlapped the *a priori* ROIs, as well as in several clusters in frontal cortex in both hemispheres (Table 3). For classification of *location-relevant* vs. *order-relevant* trials, the searchlight analysis also identified clusters in areas that overlapped the *a priori* ROIs as well as superior temporal gyrus and insula (Table 4).

**Table 3.** Regions identified by searchlight MVPA to discriminate context-relevant from *context-irrelevant* trial types (see Supplementary Figure 2).

| Region | Hemisphere of cluster | Peak voxel MNI coordinates | Number of voxels | Z-score at peak |
| --- | --- | --- | --- | --- |

|  | **peak** | **(x,y,z)** | **(3.5 mm)** |  |
| --- | --- | --- | --- | --- |
| Middle frontal gyrus | Right | -43.2, -15.8, 48.2 | 359 | 4.994 |
| Inferior parietal lobule; Superior parietal lobule | Right | -29.2, 47.2, 48.2 | 197 | 4.697 |
| Inferior frontal gyrus | Left | 51.2, -33.2, 16.8 | 141 | 4.323 |
| Superior frontal gyrus; Superior middle gyrus | Right | -15.2, -33.2, 48.2 | 113 | 4.649 |
| Middle occipital gyrus | Left | 33.8, 68.2, 27.2 | 85 | 4.361 |
| Middle temporal gyrus | Left | 65.2, 33.2, -0.8 | 72 | 4.610 |
| Inferior temporal gyrus; Middle temporal gyrus | Right | -46.8, 68.2, -4.2 | 52 | 4.199 |
| Cuneus; Superior parietal lobule; Precuneus; Superior occipital gyrus | Left | 12.8, 71.8, 37.8 | 49 | 4.985 |

**Table 4.** Regions of significant searchlight classification accuracy of *location-relevant* from *order-relevant* trial types (see Supplementary Figure 3).

| **Region** | **Hemisphere of cluster peak** | **Peak voxel MNI coordinates (x,y,z)** | **Number of voxels (3.5 mm)** | **Z-score at peak** |
| --- | --- | --- | --- | --- |
| Superior temporal gyrus; Insula | Right | -50.2, 1.8, -0.8 | 85 | 4.375 |
| Precuneus | Right | -1.2, 75.2, 58.8 | 72 | 4.641 |
| Superior parietal lobule | Left | 23.2, 57.8, 55.2 | 60 | 4.157 |
| V1 | Left | 2.2, 82.2, -7.8 | 46 | 4.899 |

Controllability of context binding (*Hypothesis 3*). Broadly, the rationale for these analyses was to explore the idea that the representation of stimulus context in VWM is under strategic control, by assessing whether (and if so, how) the strength of the representation of the location of a stimulus (i.e., its location context) varied as a function of trial type (i.e., as a function of its relevance for behavior). Strength of representation was operationalized as slope of IEM reconstruction. To provide a benchmark against which the analyses of theoretical interest (i.e., of information held in VWM) could be compared, we began by applying our procedure to the representation of the location of the probe on the screen (i.e., a perceptual representation). In the *occipital-sample* ROI, IEMs trained on the location of the sample from load-of-1 trials produced robust reconstructions of the location of the probe at TRs 9 and 10 for all three load-of-3 trial types (all bootstrapped *p*-values < .001; left panel of Figure 5A), and these effects did not differ by trial type (all *t*-statistics ≤ 1.9614, all *p*-values ≥ .07; all *BFs* ≤ 1.1881). This confirmed the prediction of *Hypothesis 3A* that the strength of the neural representation of the physical location of the probe would not vary as a function of trial type (i.e., as a function of the relevance of that information).

**Figure 5.**
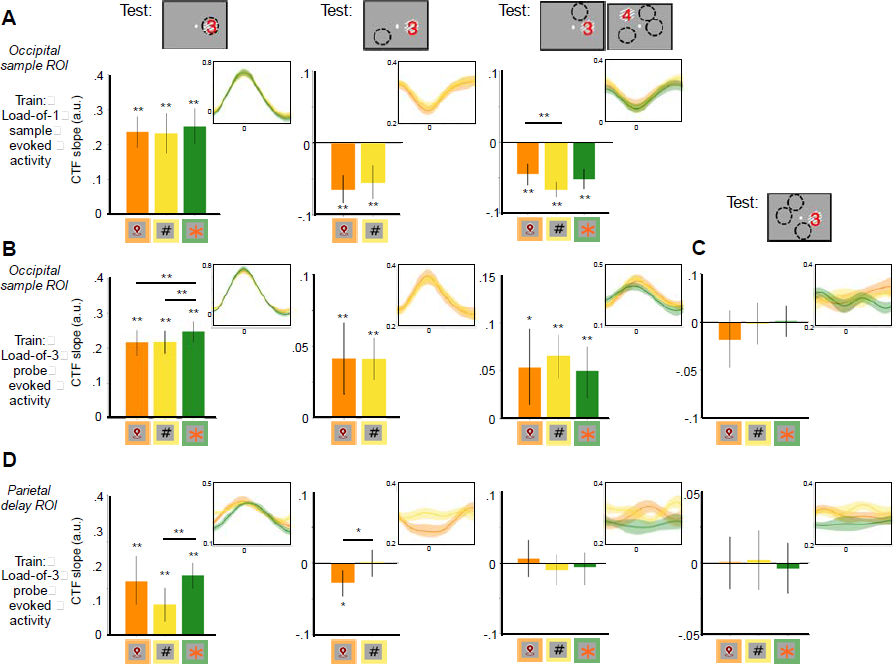
IEM reconstructions of location information from the probe-evoked signal. The illustrations of the probe display at the top of each column indicate the location (dashed circle) reconstructed by the IEM (c.f., Fig. 1): that of the item that appeared at the same location as the probe (left-hand column); that of the item referenced by the digit superimposed on the orientation patch (second from left-hand column; that of the item unreferenced by the probe (third from left-hand column); or one of the locations that was not occupied by a sample on that trial (panel C and right-hand column of panel D). Symbols along the horizontal axis of each plot indicate the trial type on which the IEM is being tested. (Note that for the second column from the left, data from context-irrelevant trials could not be used because the probe on these trials was superimposed with a digit (“4”, “5”, or “6”) that did not refer to a location at which a sample had appeared.) **(A)** *Hypothesis 3: Occipital-sample* ROI, IEMs trained on sample-evoked activity (TR 4) from load-of-1 trials: slopes of the reconstructions (CTFs circularly shifted to a common center of 0° are depicted in the insets using same color-coding as bar graphs) of the tested sample location during the load-of-3 trial type indicated by the symbol along the horizontal axis. **(B)** IEM_post hoc 2 #s 1-3_ (i.e., using models trained and tested on data from TRs 9 and 10 from load-of-3 trials), *occipital-sample* ROI. Left panel (IEM_post hoc 2 #1_): Slopes of the reconstructions with a model trained and tested on the physical location of the probe. Central panel (IEM_post hoc 2 #2_): Slopes of the reconstructions with a model trained and tested on the digit-referenced sample location. Right panel (IEM_post hoc 2 #3_): Slopes of the reconstructions with a model trained and tested on the unreferenced sample location. **(C)** IEM_post hoc 2 #4_, o*ccipital-sample* ROI: Slopes of reconstructions with a model trained and tested on locations that were not occupied by a sample on that trial. **(D)** IEM_post hoc 2 #s 1-4_*, parietal-delay* ROI: Reconstructions using the same procedures as **(B)** and **(C)**. Ribbon widths in the CTF inset plots depict +/−1 *SEM;* error bars in the bar graphs correspond to bootstrapped 95% confidence intervals. “*” and “**” above the individual bars correspond to bootstrapped *p*-values < .05 & < .01, respectively. “*”

The analyses of principal theoretical interest focused on context reinstatement: At the end of the trial is context reinstatement specific to the item being cued for the recognition judgment (*Hyp. 3B*), or is it nonspecific (i.e., is a representation of the location of all three sample stimuli reinstated; *Hyp. 3C*)? Importantly, the same encoding model was used for all of the analyses reported here. First, we report IEM reconstructions of the location of the sample that was referenced by the digit superimposed on the probe stimulus (*Hypothesis 3B*). In the *occipital-sample* ROI, IEM reconstructions of the location of the digit-referred sample were significantly negative (all bootstrapped *p*-values ≤ .0228), and they did not differ between *location-relevant* and *order-relevant* trial types (t(14) = 0.8921, *p* = .2152, *BF_10_*= 0.00004). Therefore, they were opposite in sign relative to, and significantly different from, the reconstructions of the physical location of the probe (i.e., results for *Hyp. 3A*; all *t*-statistics ≥ 5.3097, all *p*-values < .001; all *BFs* ≥ 231.3).

Next, we turn to IEM reconstructions of the location of the sample that did not correspond to either the location-on-the-screen of the probe or the probe’s digit (*Hypothesis 3C*). For this item, the slopes of the reconstructions of its location were, again, significantly negative (all bootstrapped *p*-values < .001). Additionally, they were significantly more negative on *order-relevant* trials than on *location-relevant* trials (*t*(14) = 2.5633, *p =* .0225, *BF_10_* = 2.8798), with no other differences observed (middle and right panels of Figure 5A). We found the same qualitative pattern of results when we repeated these analyses but trained the model on the probe-evoked signal (TRs 9 and 10) from the load-of-3 trial types (results not shown). The negative reconstructions generated by these analyses are almost surely due to the fact that activity in the *occipital-sample ROI* during the probe/response period of the trial was dominated by the visual drive of the probe stimulus on the screen. These outcomes, although unexpected and ill-suited for testing the encoding of locations other than that of the probe (*Hypotheses 3B and 3C)*, gave rise to ideas for alternative approaches, which we present here as post hoc analyses.

*Post hoc analyses. Post hoc analysis 1* revealed significant positive reconstruction of the location of the to-be probed sample item during the late delay period (TRs 6 and 7) in the *occipital-sample ROI* for *location-relevant* trials only (both bootstrapped *p*-values ≤ .0104; all other bootstrapped *p-*values ≥ .0516 for reconstructions for other trial types and for reconstructions in the *parietal-delay* ROI). Direct comparison of the location-relevant and context-irrelevant trial type reconstruction strengths in the *occipital-sample* ROI at these TRs revealed a significant difference between the two conditions (both t-statistics ≥ 2.2498, both p-values ≤ .041; all BFs ≥ 1.7906).

*Post hoc analysis 2* addressed the question of context reinstatement at the time of the probe by training and testing each of four IEMs on data from TRs 9 and 10, each labeled according to the location of a different sample item. The results in the *occipital-sample* ROI produced significantly positive reconstructions of the location of the probe on the screen (on all three trial types; IEM_post hoc 2 #1_), of the location of the digit-cued item (on both *location-relevant* and *order-relevant* trials; IEM_post hoc 2 #2_), and of the location of the uncued item(s) (the uncued item on *location-relevant* and *order-relevant* trials, and all three locations on *context-irrelevant* trials; IEM_post hoc 2 #3_) (all bootstrapped *p*-values < .02; Figure 5B). The interpretability of these results was reinforced by the failure of IEM_post hoc 2 #4_ to reconstruct, at TRs 9 and 10, the locations that had not been occupied by a sample (all bootstrapped *p*-values ≥ .4364; Figure 5C). Together, the results from *post hoc analysis 2* suggest that, at test, and in occipital cortex, the locations of all sample items from that trial were reactivated, regardless of whether or not they corresponded to the item being probed.

In the *parietal-delay* ROI, results for IEM_post hoc 2 #1_ revealed significant positive reconstruction of the location-on-the-screen of the probe for all three trial types (all bootstrapped *p*-values < .002), with the slope of the reconstruction during *order-relevant* trials numerically lower than during the other two trial types, and significantly so for *context-irrelevant* trials (*t*(14) = 2.6459, *p =* .0192; *BF_10_* = 3.2771). Results for IEM_post hoc 2 #2_ (trained and tested on the location of the item referenced by the superimposed digit in the probe stimulus) revealed a significant *negative* reconstruction of the location of the digit-referenced item on *location-relevant* trials (bootstrapped *p*-value = .0048), an effect that differed significantly from its reconstruction on *order-relevant* trials (*t*(14) = 2.2801, *p =* .0388; *BF_10_* = 1.8725). For IEM_post hoc 2 #3_ (trained and tested on the location of the item that was not cued by the probe stimulus) and IEM_post hoc 2 #4_ (trained and tested on the locations not occupied by a sample), no significant reconstructions were observed (all bootstrapped p-values ≥ 0.4336; Figure 5D). Therefore, the patterns from the *parietal-delay* ROI differed markedly from those in the *occipital-sample* ROI, suggesting that the former may have played a role in deemphasizing irrelevant information during test (i.e., the location of the order-referred item on *location*-*relevant* trials (Figure 5D, second from left), and the location of the location-referred item on *order-relevant* trials (Figure 5D, left-hand column).^1^

Because the interpretation of IEM reconstructions with negative slopes can be equivocal, we carried out simulations to assess whether we could replicate the empirical findings from the parietal delay ROI (Figure 5D; c.f., Adam & Serences, 2021). When all three were given equal weight, the simulations generated robust positive reconstructions of the location of each of the three sample items. When the activation of the digit-referenced sample item was downweighted, however, the simulation produced a reconstruction with a negative slope (see Supplementary Materials for more detail). These simulation results are therefore consistent with the interpretation that the pattern of IEM reconstructions observed in the parietal-delay ROI cortex may reflect the filtering of task-irrelevant stimulus information.

## Discussion

Context plays a critical role in working memory when situations require memory for where and/or when an item was encountered. Recent research has identified sensitivity to context-binding demands in the delay-period activity of IPS (Gosseries, Yu, et al. 2018; Cai et al. 2019; Cai et al. 2020), indicating a role above and beyond that of item representation (Todd & Marois 2004, Todd & Marois 2005; Xu & Chun 2006). The results presented here replicate this evidence for context-binding sensitivity of IPS, and extend it to a task in which all sample items were drawn from the same category, sample displays were identical across trials, and only the informational domain of trial-critical context varied on a trial-by-trial basis. This and several other aspects of the present results, to be considered below, demonstrate important regional differences in the processing of context in VWM. They suggest that whereas occipital cortex supports the representation of stimulus context in a manner that is task invariant and perhaps automatic, IPS may support the strategic up- and down-weighting of this contextual information to effect the selective filtering of information held in VWM. This latter profile is consistent with the function of a priority map (c.f., Zelinsky & Bisley 2015).

One piece of evidence for a functional dissociation of IPS from occipital cortex came from the first question motivating this experiment: the confirmation of *Hypothesis 1*’s predictions of differential patterns of sensitivity to context-binding demands. Across trials that presented identical displays of three to-be-remembered samples, only in IPS was delay-period activity for context-relevant than *context-irrelevant* trials. Exploratory whole-brain univariate analyses revealed differences in delay-period activity for context-relevant and *context-irrelevant* trials in several clusters in parietal and frontal cortex, as well as in the cerebellum and basal ganglia, with no differences observed for *location-relevant* and *order-relevant* trials in any region, including IPS.

The analyses motivated by our second question *–* addressing the informational domain of stimulus context -- also revealed differences between occipital cortex and PPC. In occipital cortex MVPA decoding of trial-specific context information (i.e., ‘*what kind of trial is it:* location, order, *or* irrelevant*?*’) was strongest for the instructional cue and stimulus encoding, then dropped to near-chance levels for the remainder of the trial. In PPC, in contrast it grew steadily and peaked at the time of the memory-guided response. In addition to occipital cortex and PPC, whole brain analysis identified several additional regions whose activity discriminated *location-relevant* from *order-relevant* trials. We note, however, that results from these analyses cannot support strong interpretation of a region’s possible selectivity for one domain versus another, because they cannot discriminate, for example, a region specialized for processing spatial context from one involved in the processing of spatial context and ordinal context.

The results addressing our third question -- whether the processing of stimulus context can be susceptible to cognitive control – again highlight marked differences in the VWM functions of occipital cortex vs. PPC. In occipital cortex, the fact that the locations of each of the three samples were actively represented during the probe epoch, on all three trial types, provides evidence for the automatic reinstatement of the location context of all items currently in VWM, regardless of the relevance of this information for selecting the item cued by the probe. This evidence for an automatic reinstatement of location context is consistent with previous evidence for the incidental encoding of location information, regardless of its task relevance (e.g., Ellis 1990; Treisman & Zhang 2006; Clark et al. 2012; Kondo & Saiki 2012; Foster et al. 2017; Cai et al. 2019; Heuer & Rolfs 2021)^2,3^. In addition to the putatively automatic activation of location that accompanies probe onset, the data also showed evidence that this information can be selectively, perhaps strategically, activated in advance of probe onset, on trials when it will be needed for the recognition decision. Importantly, the failure to find these effects with IEMs trained on sample location from load-of-1 trials suggests that the neural code representing reinstated location context late in the delay period differs from that representing the sensory representation of the perceived location of an item on the screen. Alternatively, it could be an indication that the binding of location context to stimulus information on load-of-3 trials is not purely automatic, but is being carried out strategically to disambiguate the three items (i.e., in a way that is not needed for a single item).

In parietal cortex, the results of IEM reconstructions of location context were markedly different (compare Fig. 5B vs. **5D**), and were more consistent with a role in filtering or weighting information according to its relevance for behavior (c.f., Zelinsky & Bisley 2015). In particular, the strength of representation of the location of the probe was lower on *order-relevant* trials (when it was not relevant), and that of the location of the digit-referred stimulus was lower on *location-relevant* trials. Indeed, the IEM reconstruction of the location of the digit-referred item had a negative slope on *location-relevant* trials, a pattern opposite of its reconstruction on these same trials in occipital cortex. This specific result suggests a function of deemphasizing an item whose representation might otherwise interfere with the recognition decision. Simulations indicated that these results can be produced by the selective down-weighting of this information.^4^ A limitation to acknowledge here is that our study was not designed to provide evidence for active selection, as might be expected from an IPS-based priority map (e.g., Bisley & Goldberg 2010; Jerde et al. 2012; Bisley & Mirpour, 2019).

When we consider behavioral performance, it seems likely that part of the superiority of *location-relevant* trials relative to the other two trial types is a ‘same-position’ advantage (Hollingworth 2007; Sapkota et al. 2011). Additionally, however, the neural evidence for the probe-triggered activation of the location context of all items during *order-relevant* and *context-irrelevant* trials suggests the possibility that performance on these trials may also have suffered from interference from this trial-irrelevant information. Additional research with a procedure similar to the one used here, but testing recall instead of recognition, will be needed to explore whether the trial-irrelevant representation of location context may degrade performance on *order-relevant* and *context-irrelevant* trials.

### Citation diversity statement

To promote transparency surrounding citation practice (Dworkin et al. 2019; Zurn et al. 2020) and mitigate biases leading to under-citation of work led by women relative to other papers in demonstrated across several scientific domains (e.g., Dworkin et al. 2019; Maliniak et al. 2013; Caplar et al. 2017; Fulvio et al. 2021), we proactively aimed to include references that reflect the diversity of the field and quantified the gender breakdown of citations in this article according to the first names of the first and last authors using the Gender Citation Balance Index web tool (https://postlab.psych.wisc.edu/gcbialyzer/) with manual correction as needed. This article contains 51.4% man/man, 8.1% man/woman, 32.4% woman/man and 8.1% woman/woman citations. For comparison, proportions estimated from articles in five prominent neuroscience journals (as reported in Dworkin et al. 2019) are 58.4% man/man, 9.4% man/woman, 25.5% woman/man and 6.7% woman/woman. Note that the estimates may not always reflect gender identity and do not account for intersex, non-binary, or transgender individuals.

## Acknowledgements

We thank Joshua Chung for assistance with subject recruitment and behavioral screening.

## Footnotes

1 For completeness, we report here the results of additional planned analyses related to *Hypothesis 3*. For *Hyp. 3(ii)*, we were unable to reconstruct the location of any sample item during the late delay period, in either the *occipital-sample* or the *parietal-delay* ROI (all bootstrapped *p*-values ≥ .0744). For *Hyp. 3(iii)*, we were unable to reconstruct orientation in any of the conditions at any time point. The failure to reconstruct orientations in the current experiment might be due to the large number of orientations presented on each trial and/or to the peripheral presentation of orientations.

2 We note, however, that our design leaves open the possibility that other factors may also have accounted for the encoding of location information on *order-relevant* and *context-irrelevant* trials. One is that these trials were intermixed with location-relevant trials; another is that location information could have been employed strategically to help with order memory, such as by representing the series of sample items as having appeared along a path through space.

3 We further note that one limitation of our study is that the lack of an a priori model of how sequential order is represented in the brain, particularly with a code that would be discriminable with fMRI. This prevented us from carrying out analyses comparable to the IEM analyses exploring the representation of location context. Thus, an open question for future research is whether it might be possible to find neural evidence that ordinal context may also be obligatorily encoded in VWM (Heuer & Rolfs 2021), and similarly flexibly prioritized according to task-specific demands.

4 (In other research, similar patterns of "negative" IEM reconstruction have been associated with items that are either to be deprioritized (Yu, Teng, & Postle 2020; Wan et al. 2020; Wan et al. 2022), or dropped (Lorenc et al. 2020) from VWM.)

## References

Adam, K.C. & Serences, J.T. (2021). History Modulates Early Sensory Processing of Salient Distractors. Journal of Neuroscience, 41(38), 8007–8022. https://doi.org/10.1523/JNEUROSCI.3099-20.2021

Bettencourt, K. C., & Xu, Y. (2016). Decoding the content of visual short-term memory under distraction in occipital and parietal areas. Nature neuroscience, 19(1), 150–157. https://doi.org/10.1038/nn.4174

Bisley, J. W., & Goldberg, M. E. (2010). Attention, intention, and priority in the parietal lobe. Annual review of neuroscience, 33, 1–21. https://doi.org/10.1146/annurev-neuro-060909-152823

Bisley, J. W., & Mirpour, K. (2019). The neural instantiation of a priority map. Current opinion in psychology, 29, 108–112. https://doi.org/10.1016/j.copsyc.2019.01.002

Brainard, D. H., & Vision, S. (1997). The psychophysics toolbox. Spatial vision, 10(4), 433–436. https://doi.org/10.1163/156856897x00357

Cai, Y., Sheldon, A. D., Yu, Q., & Postle, B. R. (2019). Overlapping and distinct contributions of stimulus location and of spatial context to nonspatial visual short-term memory. Journal of neurophysiology, 121(4), 1222–1231. https://doi.org/10.1152/jn.00062.2019

Cai, Y., Fulvio, J. M., Yu, Q., Sheldon, A. D., & Postle, B. R. (2020). The role of location-context binding in nonspatial visual working memory. Eneuro, 7(6). https://doi.org/10.1523/ENEURO.0430-20.2020

Caplar, N., Tacchella, S., & Birrer, S. (2017). Quantitative evaluation of gender bias in astronomical publications from citation counts. Nature astronomy, 1(6), 1–5. https://doi.org/10.1038/s41550-017-0141

Clark, K. L., Noudoost, B., & Moore, T. (2012). Persistent spatial information in the frontal eye field during object-based short-term memory. Journal of Neuroscience, 32(32), 10907–10914. https://doi.org/10.1523/JNEUROSCI.1450-12.2012

Cox, R. W. (1996). AFNI: software for analysis and visualization of functional magnetic resonance neuroimages. Computers and Biomedical research, 29(3), 162–173. https://doi.org/10.1006/cbmr.1996.0014

Dworkin, J. D., Linn, K. A., Teich, E. G., Zurn, P., Shinohara, R. T., & Bassett, D. S. (2020). The extent and drivers of gender imbalance in neuroscience reference lists. Nature neuroscience, 23(8), 918–926. https://doi.org/10.1038/s41593-020-0658-y

Ellis, N. R. (1990). Is memory for spatial location automatically encoded?. Memory & Cognition, 18(6), 584–592. https://doi.org/10.3758/BF03197101

Foster, J. J., Bsales, E. M., Jaffe, R. J, & Awh, E. (2017). Alpha-band activity reveals spontaneous representations of spatial position in visual working memory. Current Biology, 27, 3216–3223.e6, 2017. https://doi.org/10.1016/j.cub.2017.09.031

Fulvio, J. M., Akinnola, I., & Postle, B. R. (2021). Gender (im) balance in citation practices in cognitive neuroscience. Journal of Cognitive Neuroscience, 33(1), 3–7. https://doi.org/10.1162/jocn_a_01643

Gosseries, O., Yu, Q., LaRocque, J. J., Starrett, M. J., Rose, N. S., Cowan, N., & Postle, B. R. (2018). Parietal-occipital interactions underlying control-and representation-related processes in working memory for nonspatial visual features. Journal of Neuroscience, 38(18), 4357–4366. https://doi.org/10.1523/JNEUROSCI.2747-17.2018

Hebart, M. N., Görgen, K., & Haynes, J.-D. (2015). The Decoding Toolbox (TDT): a versatile software package for multivariate analyses of functional imaging data. Frontiers in Neuroinformatics, 8(January), 1–18. https://doi.org/10.3389/fninf.2014.00088

Heuer, A., & Rolfs, M. (2021). Incidental encoding of visual information in temporal reference frames in working memory. Cognition, 207, 104526. https://doi.org/10.1016/j.cognition.2020.104526

Hollingworth, A. (2007). Object-position binding in visual memory for natural scenes and object arrays. Journal of Experimental Psychology: Human Perception and Performance, 33(1), 31. https://doi.org/10.1037/0096-1523.33.1.31

Jerde, T. A., Merriam, E. P., Riggall, A. C., Hedges, J. H., & Curtis, C. E. (2012). Prioritized maps of space in human frontoparietal cortex. Journal of Neuroscience, 32(48), 17382–17390. https://doi.org/10.1523/JNEUROSCI.3810-12.2012

Kleiner, M., Brainard, D., & Pelli, D. (2007). What’s new in Psychtoolbox-3?.

Kondo, A., & Saiki, J. (2012). Feature-specific encoding flexibility in visual working memory. PloS one, 7(12), e50962. https://doi.org/10.1371/journal.pone.0050962

Lorenc, E. S., Vandenbroucke, A. R., Nee, D. E., de Lange, F. P., & D’Esposito, M. (2020). Dissociable neural mechanisms underlie currently-relevant, future-relevant, and discarded working memory representations. Scientific reports, 10(1), 1–17. https://doi.org/10.1038/s41598-020-67634-x

Maliniak, D., Powers, R., & Walter, B. F. (2013). The gender citation gap in international relations. International Organization, 67(4), 889–922. https://doi.org/10.1017/S0020818313000209

Pelli, D. G., & Vision, S. (1997). The VideoToolbox software for visual psychophysics: Transforming numbers into movies. Spatial vision, 10, 437–442. https://doi.org/10.1163/156856897x00366

Sapkota, R. P., Pardhan, S., & van der Linde, I. (2011). Object—Position Binding in Visual Short-Term Memory for Sequentially Presented Unfamiliar Stimuli. Perception, 40(5), 538–548. https://doi.org/10.1068/p6899

Sprague, T.C., Adam, K.C., Foster, J.J., Rahmati, M., Sutterer, D.W., Vo, V.A. (2018). Inverted encoding models assay population-level stimulus representations, not single-unit neural tuning. eNeuro 5:ENEURO.0098-18.2018. https://doi.org/10.1523/ENEURO.0098-18.2018

Sprague, T.C., Boynton, G.M., Serences, J.T. (2019). The importance of considering model choices when interpreting results in computational neuroimaging. eNeuro 6:ENEURO.0196-19.2019. https://doi.org/10.1523/ENEURO.0196-19.2019

Todd, J. J., & Marois, R. (2004). Capacity limit of visual short-term memory in human posterior parietal cortex. Nature, 428(6984), 751–754. https://doi.org/10.1038/nature02466

Todd, J. J., & Marois, R. (2005). Posterior parietal cortex activity predicts individual differences in visual short-term memory capacity. Cognitive, Affective, & Behavioral Neuroscience, 5(2), 144–155. https://doi.org/10.3758/CABN.5.2.144

Treisman, A., & Zhang, W. (2006). Location and binding in visual working memory. Memory & cognition, 34(8), 1704–1719. https://doi.org/10.3758/BF03195932

van Loon, A. M., Olmos-Solis, K., Fahrenfort, J. J., & Olivers, C. N. (2018). Current and future goals are represented in opposite patterns in object-selective cortex. ELife, 7: e38677. https://doi.org/10.7554/eLife.38677

Wan, Q., Cai, Y., Samaha, J., & Postle, B. R. (2020). Tracking stimulus representation across a 2-back visual working memory task. Royal Society open science, 7(8), 190228. https://doi.org/10.1098/rsos.190228

Wan, Q., Menendez, J. A., & Postle, B. R. (2022). Priority-based transformations of stimulus representation in visual working memory. PLOS Computational Biology, 18(6), e1009062. https://doi.org/10.1371/journal.pcbi.1009062

Xu, Y. (2017). Reevaluating the sensory account of visual working memory storage. Trends in Cognitive Sciences, 21(10), 794–815. https://doi.org/10.1016/j.tics.2017.06.013

Xu, Y., & Chun, M. M. (2006). Dissociable neural mechanisms supporting visual short-term memory for objects. Nature, 440(7080), 91–95. https://doi.org/10.1038/nature04262

Yu, Q., Teng, C., & Postle, B. R. (2020). Different states of priority recruit different neural representations in visual working memory. PLoS biology, 18(6), e3000769. https://doi.org/10.1371/journal.pbio.3000769

Zelinsky, G. J., & Bisley, J. W. (2015). The what, where, and why of priority maps and their interactions with visual working memory. Annals of the new York Academy of Sciences, 1339(1), 154–164. https://doi.org/10.1111/nyas.12606

Zurn, P., Bassett, D. S., & Rust, N. C. (2020). The citation diversity statement: a practice of transparency, a way of life. Trends in Cognitive Sciences, 24(9), 669–672. https://doi.org/10.1016/j.tics.2020.06.009

