## Supplementary for "Strategic control of location and ordinal context in visual working memory"

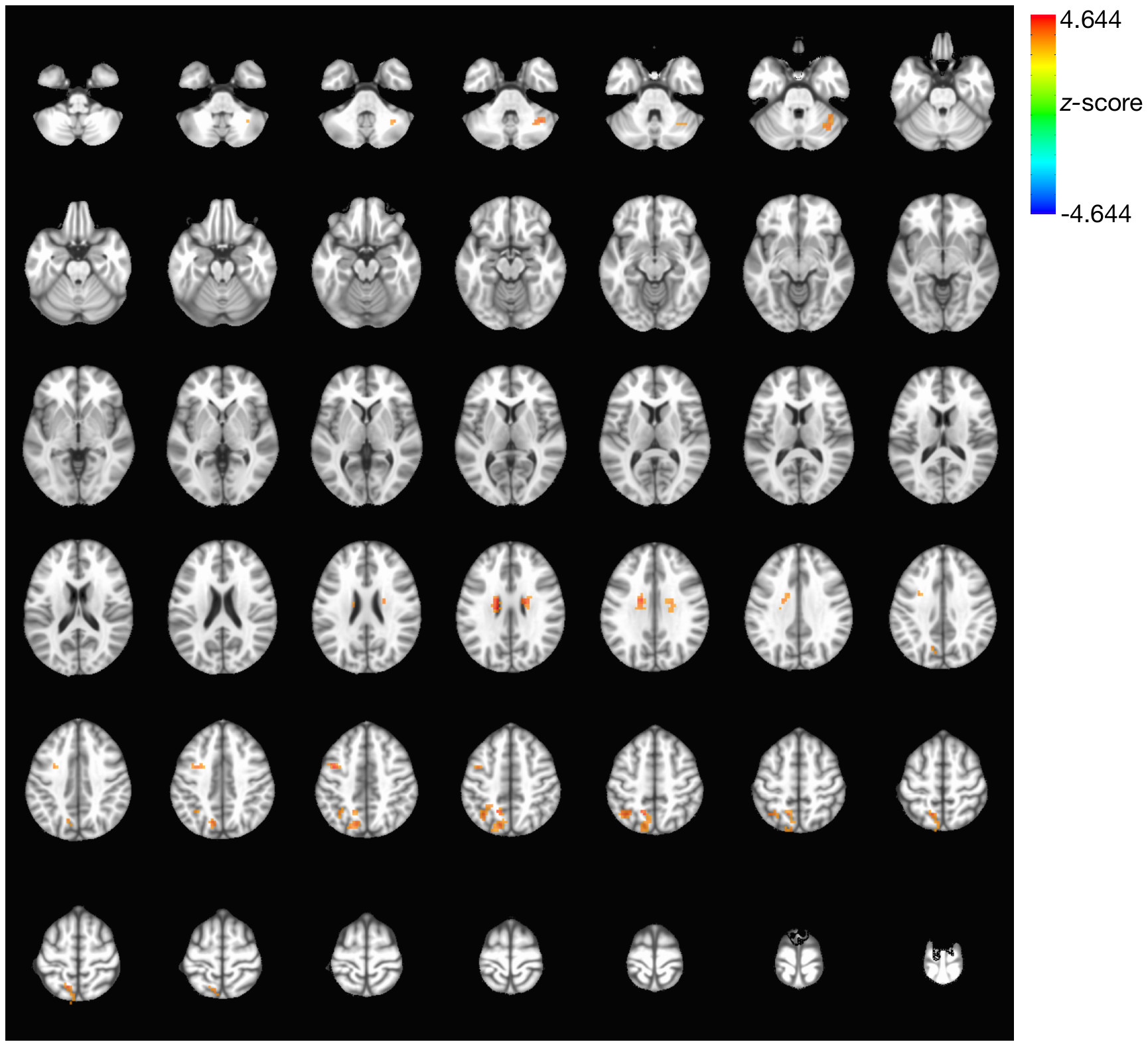

**Supplementary Figure 1.** Results from whole-brain contrast of delay-period activity during context-relevant vs. *context-irrelevant* trials, displayed in neurological convention on axial anatomical images from z = -45 (upper left) to z = 78 (lower right).

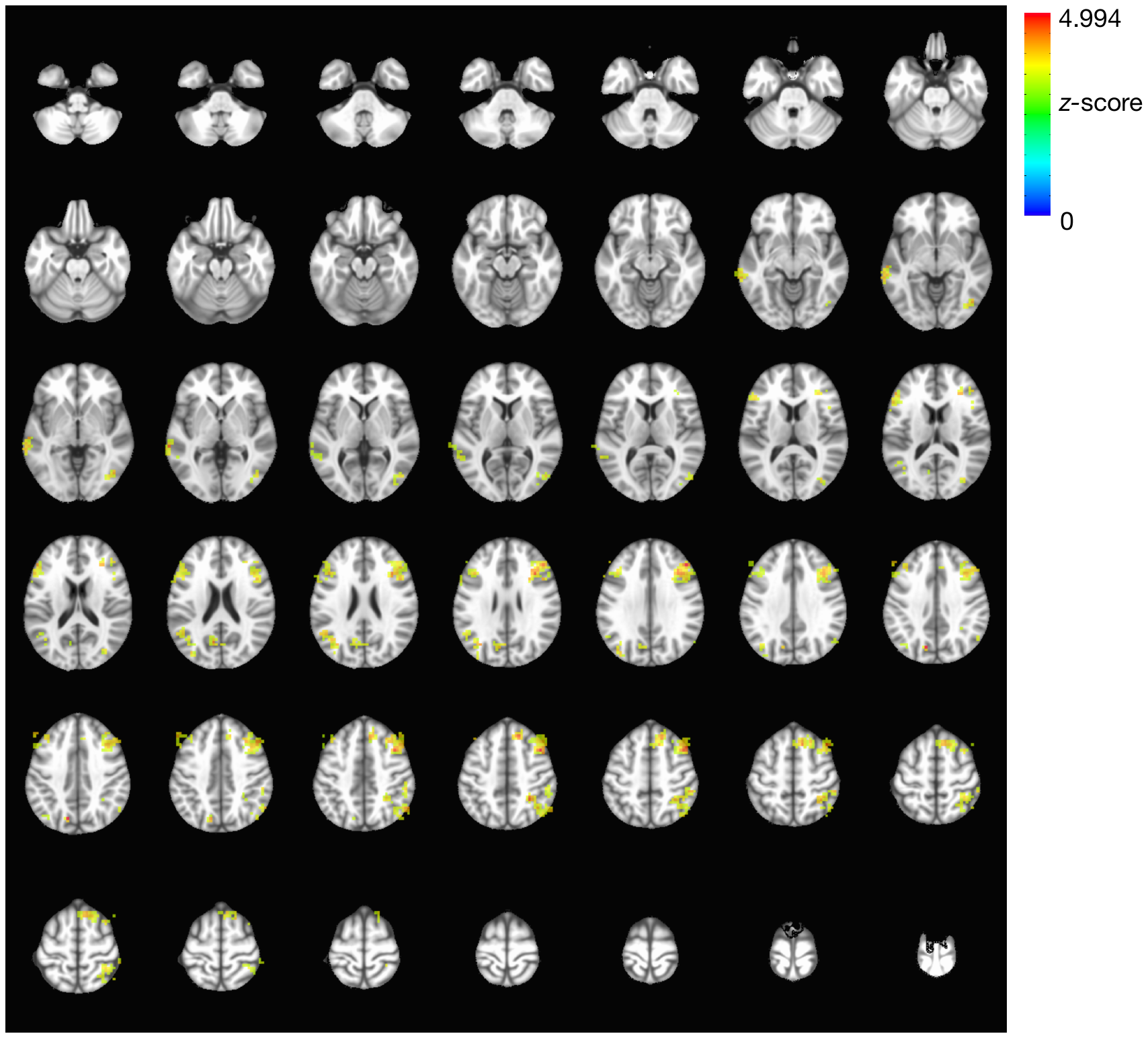

**Supplementary Figure 2.** Clusters identified in a searchlight MVPA to discriminate context-relevant from *context-irrelevant* trial types. Same display conventions as Supp. Fig. 1.

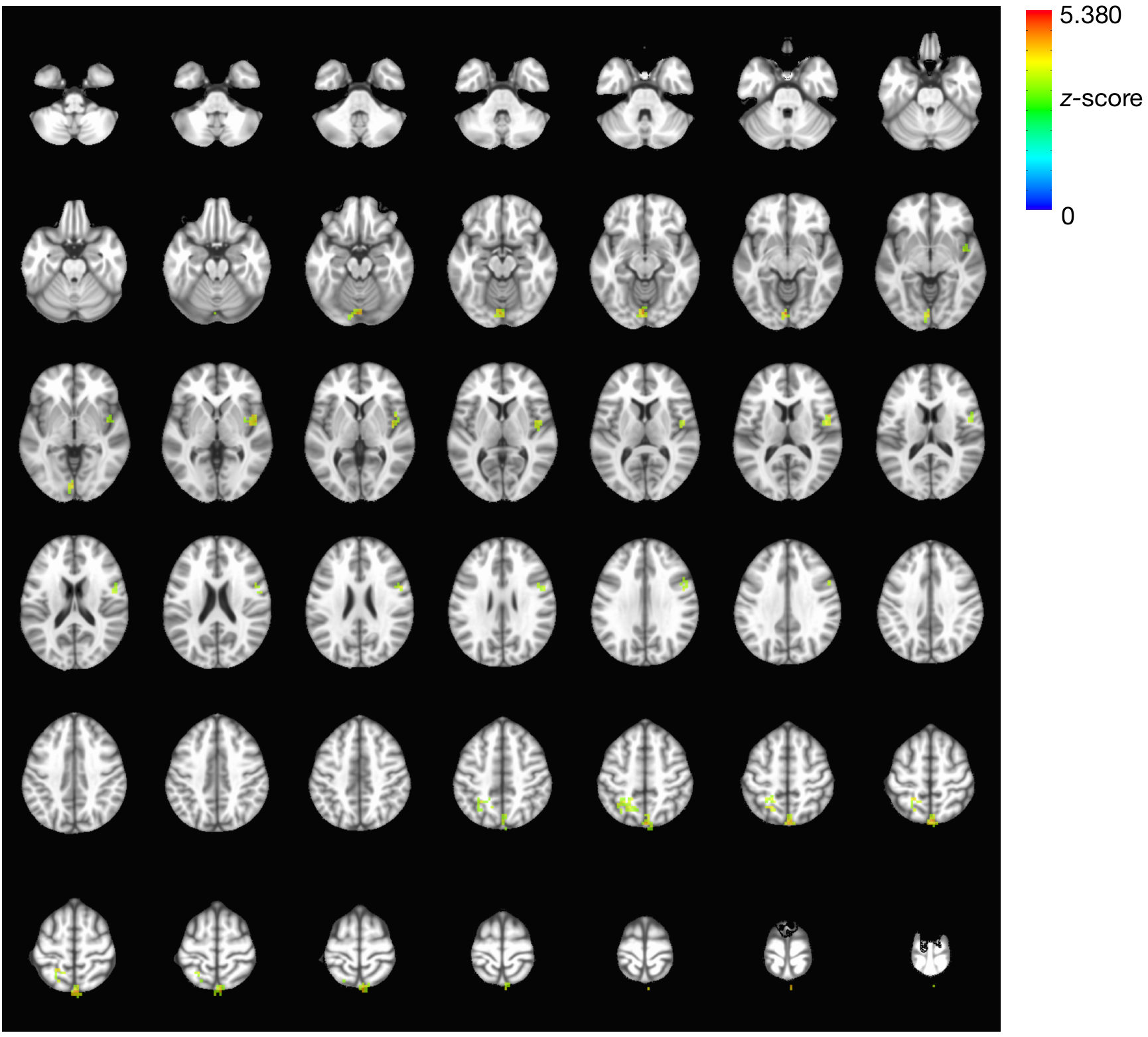

**Supplementary Figure 3.** Clusters identified in a searchlight MVPA to discriminate location-relevant from order-relevant trial types. Same display conventions as Supp. Fig. 1.

**IEM Simulations**

Following Adam & Serences (2021), we used data from the Load-of-1 trials to generate predictions for observed model responses to a 3-item memory set. For each subject, we trained the model on the Load-of-1 sample TR (TR 4) BOLD signal data from the *occipital sample* ROI, and recorded the model response derived from the probe period (TR 9) for each of the six sample locations. We then computed the average CTF of the reconstruction of Load-of-1 probe-evoked signal (TR9) across sample locations and subjects for use as the ‘canonical’ reconstruction in the simulations.

The simulation step consisted of replicating the canonical reconstruction three times (once at each of the three sample item locations). Depending on the sample item location we aimed to reconstruct, we shifted that item’s reconstruction to be centered on 0 and shifted the reconstruction of the remaining items relative to that. For example, if we aimed to reconstruct the physical location of the probe on the screen, we aligned the probe’s reconstruction to 0 and the reconstructions of the two remaining sample items were shifted by the distance between their locations and the probe’s location. We then took the average of the three reconstructions to generate a single model prediction for each trial – this comprised the ‘equal weight’ simulation as the reconstructions of all three items contributed equally to the final prediction for each trial.

Inspection of individual trial predictions revealed a variety of patterns. This is a consequence of the fact that the three sample item locations were randomly selected on each trial (but were never overlapping) and hence varied in their distances from each other. **Figure S4** shows several examples, including trials with three distinct peaks, trials with single 0-centered peaks, and trials with single non-0-centered peaks, trials with multiple less well-defined peaks.

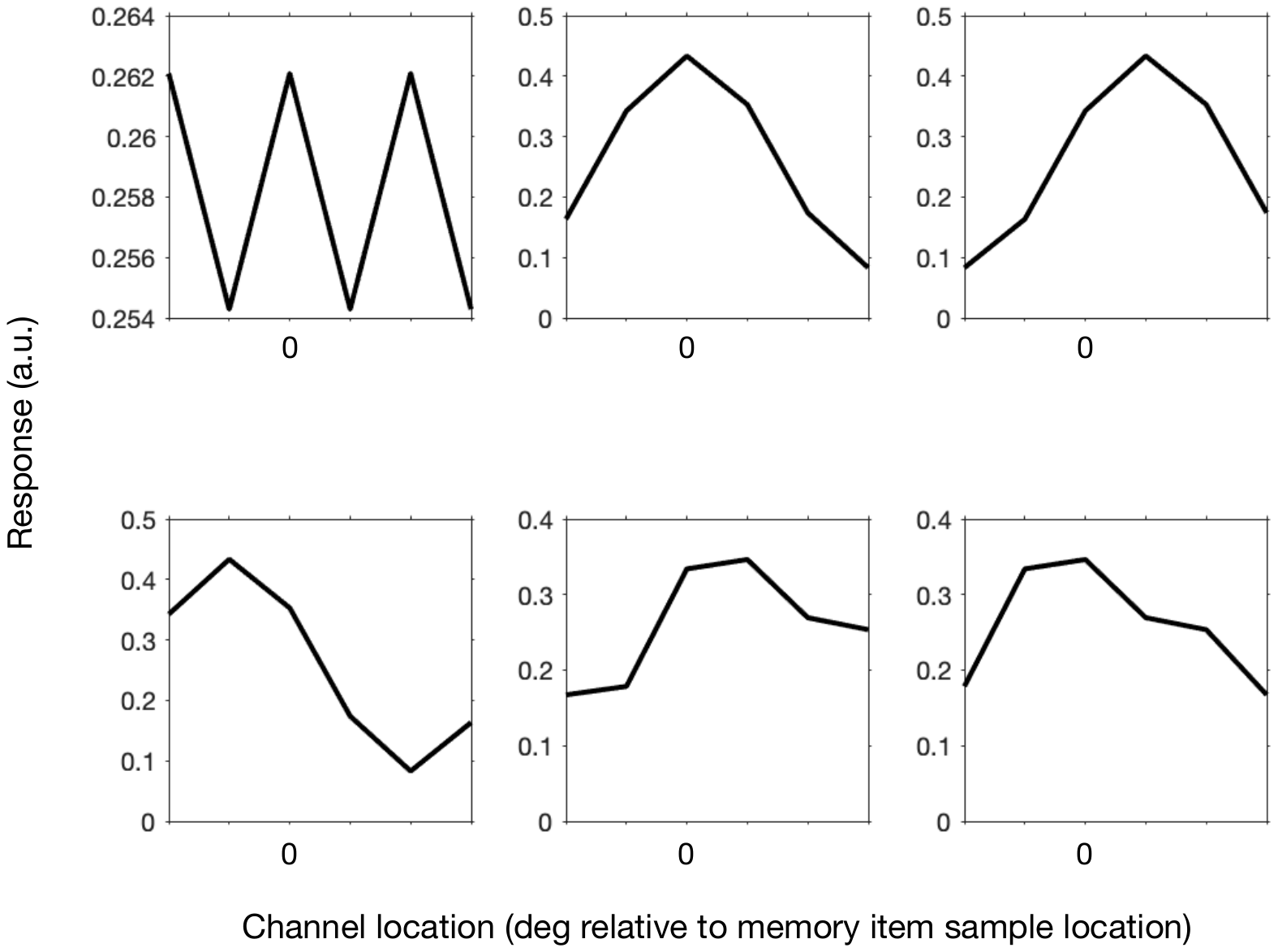

**Figure S4.** Example individual-trial model predictions (a.k.a., “IEM reconstructions)”for Load-of-3 trials when all three memory items are given the same weight at probe (TR 9).

To obtain an averaged model response for the reconstruction of the probe’s physical location, we averaged over all trials when the response was 0-center aligned to the probe’s physical location. Likewise, we obtained an average model response for the reconstruction of the digit-referenced sample location and the unreferenced sample location by averaging over all trials when the responses were 0-center aligned to those locations. The top row of **Figure S5** shows that when the three memory items are given the same weight at probe, a positive, equivalent reconstruction is predicted for all three memory item locations.

We then repeated the simulation step but varied the strength of the simulated response to each item to explore the effects on the reconstructions. We first simulated a downweighting or suppression of the non-probe locations by 50%. This was achieved by multiplying the canonical reconstruction by .5 before shifting to the digit-referenced item location and the unreferenced item location. The middle row of **Figure S5** shows that when the non-probe locations are downweighted at probe, positive reconstructions are predicted for all three memory item locations, but these reconstructions have less strength than that of the probe location reconstruction.

Finally, we repeated the downweighting analysis by simulating a suppression of 90%, which was achieved by multiplying the canonical reconstruction by .1 for the two non-probe locations. The bottom row of **Figure S5** shows that in this case, *negative* reconstructions are predicted for the non-probe locations.

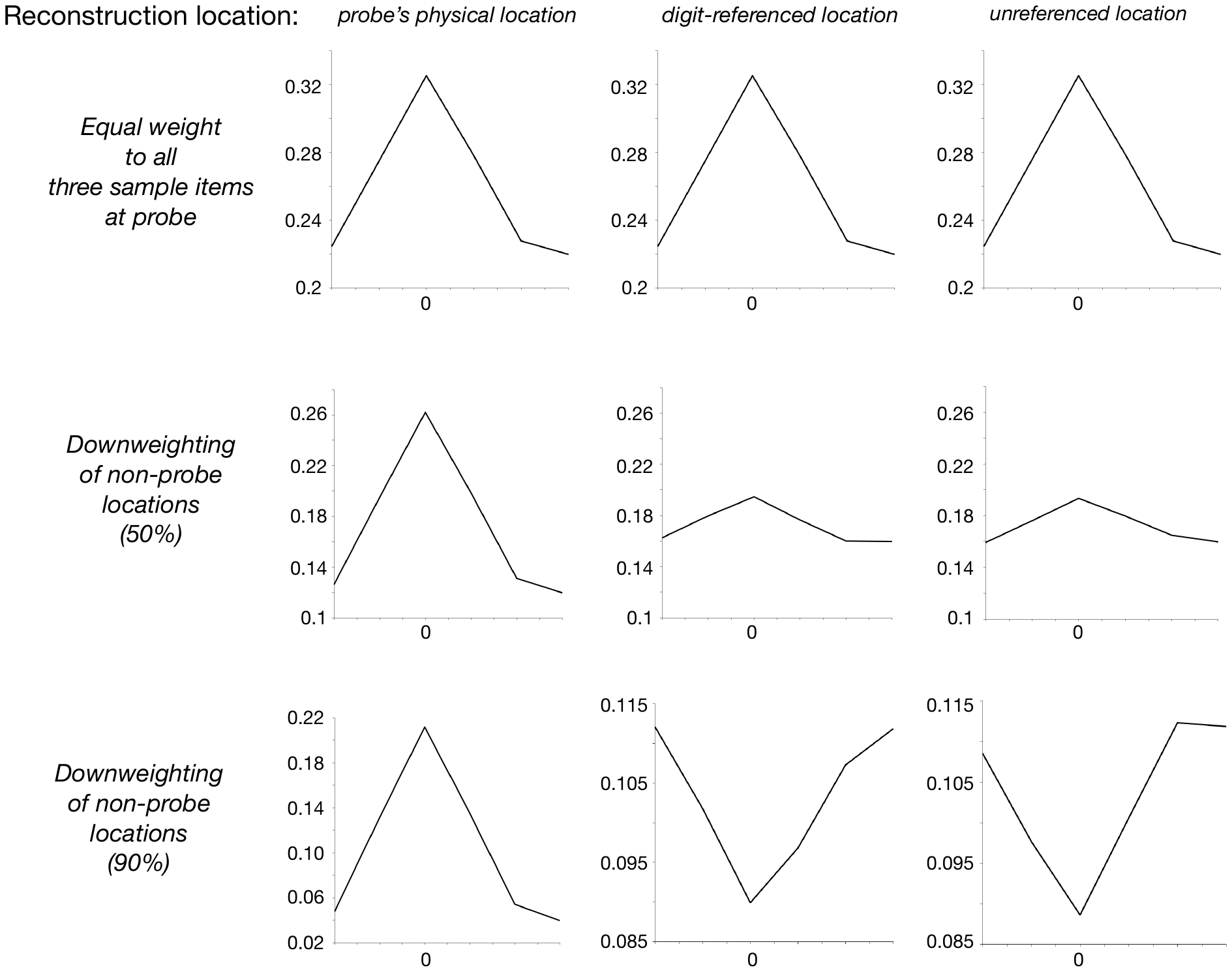

**Figure S5.** Simulated Load-of-3 probe period model reconstructions when aligned to the probe’s physical location (left column), the location of the digit-referenced sample item (middle column), and the location of the remaining unreferenced sample item (right column) when all three items are given equal weight (upper row), when the two non-probe location reconstructions are downweighted by 50% (middle row), and when the two non-probe location reconstructions are downweighted by 90%.

Lastly, we carried out a final analysis in which we simulated the model reconstructions in the location-relevant condition and order-relevant condition separately. In the location-relevant condition, we simulated *both* an enhancement of the probe’s physical location, which was achieved by multiplying the canonical reconstruction by 1.5 as well as a suppression of the digit-referenced sample’s location of 90% as above. In the order-relevant condition, we simulated a suppression of the digit-referenced sample’s location of 90%, leaving the strength of the probe’s physical location reconstruction unchanged. As shown in **Figure S6,** the results replicate the patterns observed in parietal cortex (**Figure 5D** of the main text (left and middle column)). Specifically, we found stronger positive reconstruction of the probe’s physical location in the location-relevant condition in which location is the critical context. We also see a stronger negative reconstruction of the digit-referenced sample location in the location-relevant condition, where this information is incongruent and create interference.

Taken together, the results of the simulations improve the interpretability and provide additional evidence for the interpretations of the IEM findings in the main text.

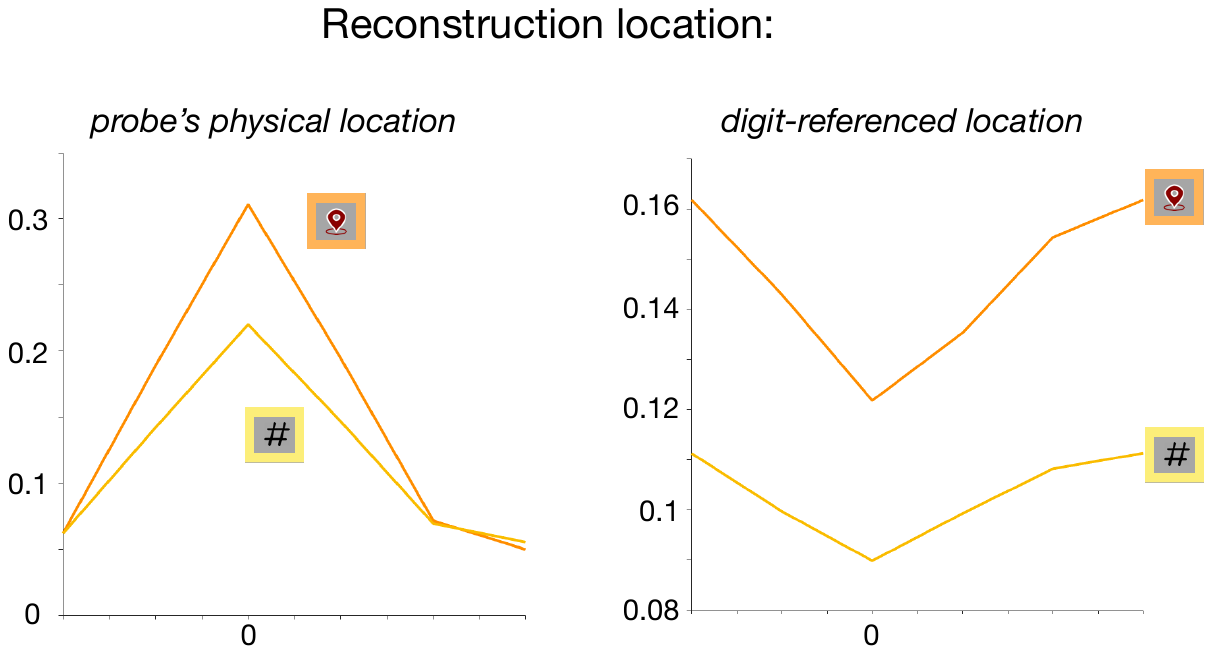

**Figure S6.** Simulated Load-of-3 probe period model reconstructions when aligned to the probe’s physical location (left column), the location of the digit-referenced sample item (middle column) for the location-relevant (location pin – orange) and order-relevant trial types (‘#’ – yellow).
